# The Taiwan Precision Medicine Initiative: A Cohort for Large-Scale Studies

**DOI:** 10.1101/2024.10.14.616932

**Authors:** Hsin-Chou Yang, Pui-Yan Kwok, Ling-Hui Li, Yi-Min Liu, Yuh-Jyh Jong, Kang-Yun Lee, Da-Wei Wang, Ming-Fang Tsai, Jenn-Hwai Yang, Chien-Hsiun Chen, Erh-Chan Yeh, Chun-yu Wei, Cathy S.-J. Fann, Yen-Tsung Huang, Chia-Wei Chen, Yi-Ju Lee, Shih-Kai Chu, Chih-hsing Ho, Cheng-Shin Yang, Yungling Leo Lee, Hung-Hsin Chen, Ming-Chih Hou, Jeng-Fong Chiou, Shun-Fa Yang, Chih-Hung Wang, Chih-Yang Huang, Kuan-Ming Chiu, Ming Chen, Fu-Tien Chiang, Sing-Lian Lee, Shiou-Sheng Chen, Wei-Jen Yao, Chih-Cheng Chien, Shih-Yao Lin, Fu-Pang Chang, Hsiang-Ling Ho, Yi-Chen Yeh, Wei-Cheng Tseng, Ming-Hwai Lin, Hsiao-Ting Chang, Ling-Ming Tseng, Wen-Yih Liang, Paul Chih-Hsueh Chen, Jen-Fan Hang, Shih-Chieh Lin, Yu-Jiun Chan, Ying-Ju Kuo, Lei-Chi Wang, Chin-Chen Pan, Yu-Cheng Hsieh, Yi-Ming Chen, Tzu-Hung Hsiao, Ching-Heng Lin, Yen-Ju Chen, I-Chieh Chen, Chien-Lin Mao, Shu-Jung Chang, Yen-Lin Chang, Yi-Ju Liao, Chih-Hung Lai, Wei-Ju Lee, Hsin Tung, Ting-Ting Yen, Hsin-Chien Yen, Chun-Ming Shih, Teh-Ying Chou, Tsan-Hon Liou, Chen-Yuan Chiang, Yih-Giun Cherng, Chih-Hwa Chen, Chao-Hua Chiu, Sung-Hui Tseng, Emily Pei-Ying Lin, Ying-Ju Chen, Hui-Ping Chuang, Tsai-Chuan Chen, Wei-Ting Huang, Joey Sin, I-Ling Liu, Yi-Chen Chen, Kuo-Kuang Chao, Yu-Min Wu, Pin-Pin Yu, Lung-Pao Chang, Kuei-Yao Yen, Li-Ching Chang, Yi-Jing Sheen, Yuan-Tsong Chen, Kamhon Kan, Hsiang-Lin Tsai, Yao-Kuang Wang, Ming-Feng Hou, Yuan-Han Yang, Chao-Hung Kuo, Wen-Jeng Wu, Jee-Fu Huang, Inn-Wen Chong, Jong-Rung Tsai, Cheng-Yu Lin, Ming-Chin Yu, Tsong-Hai Lee, Meng-Han Tsai, Yu-Che Ou, Pin-Yuan Chen, Tsung-Hui Hu, Yu-Chiau Shyu, Chih-Kuang Cheng, Yu-Jen Fang, Song-Chou Hsieh, Chien-Hung Chen, Chieh-Chang Chen, Ko-Jen Li, Chin-Hsien Lin, Hsien-Yi Chiu, Chen-Chi Wu, Chun-Yen Chen, Shi-Jye Chu, Feng-Cheng Liu, Fu-Ch Yang, Hsin-An Chang, Wei-liang Chen, Sung-Sen Yang, Yueh-feng Sung, Tso-Fu Wang, Shinn-Zong Lin, Yen-Wen Wu, Chien-Sheng Wu, Ju-Ying Jiang, Gwo-Chin Ma, Ting-Yu Chang, Juey-Jen Hwang, Kuo-Jang Kao, Chen-Fang Hung, Ting-Fang Chiu, Po-Yueh Chen, Kochung Tsui, Ming-Shiang Wu, See-Tong Pang, Shih-Ann Chen, Wei-Ming Chen, Chun-houh Chen, Wayne Huey-Herng Sheu, Jer-Yuarn Wu

**Author notes:** Corresponding authors: Ming-Shiang Wu, See-Tong Pang, Shih-Ann Chen, Wei-Ming Chen, Chun-houh Chen, Wayne Huey-Herng Sheu, Jer-Yuarn Wu. These authors contributed equally to this work (Hsin-Chou Yang, Pui-Yan Kwok, Ling-Hui Li, Yi-Min Liu, Yuh-Jyh Jong, Kang-Yun Lee, Ming-Shiang Wu, See-Tong Pang, Shih-Ann Chen, Wei-Ming Chen, Chun-houh Chen, Wayne Huey-Herng Sheu, Jer-Yuarn Wu).

## Abstract

The Taiwan Precision Medicine Initiative (TPMI), a project initiated by the Academia Sinica in collaboration with 16 major medical centers around Taiwan, has recruited 565,390 participants who consented to provide DNA samples for genetic profiling and grant access to their electronic medical records (EMR) for studies to develop precision medicine. Access to the EMR is both retrospective and prospective, allowing researchers to conduct prospective studies over time. Genetic profiling is done with population-optimized SNP arrays for the Han Chinese populations that enable genetic analyses such as genome-wide association, phenome-wide association, and polygenic risk score studies to evaluate common disease risk and pharmacogenetic response. Furthermore, the TPMI participants agree to be contacted for future research opportunities related to their genetic risks and receive personalized genetic risk profiles with health management recommendations. TPMI has established the TPMI Data Access Platform (TDAP), a central database and analysis platform that both safeguards the security of the data and facilitates academic research. The TPMI is the largest non-European cohort that merges genetic profiles with EMR in the world. With a cohort that can be followed over time, it can be utilized to validate genetic risk prediction models, conduct clinical trials to show the efficacy of risk-based health management, and optimize health policies based on genetic risks. In this report, we describe the TPMI study design, the population and genetic characteristics of the TPMI cohort, and the power it provides to conduct crucial studies in developing precision medicine on a population and personal level. As Han Chinese represent almost 20% of the world’s population, the results of TPMI studies will benefit >1.4 billion people around the world and serve as a model for developing population-based precision medicine.

## Background

Precision medicine is a global movement to improve health outcomes by tailoring medical interventions to the unique characteristics of individual patients^1,2^. For this movement to succeed, large cohorts with known disease states and rich clinical data must be built so that they can be analyzed against genetic and other factors that contribute to disease risk and treatment outcomes. Once established and validated, members of the population can match their own profiles against those from the study cohorts and identify the best medical management for their conditions. In recent years, Precision Medicine Initiatives across the world ^3^ have enriched the research landscape by producing comprehensive datasets (consisting of demographic, genetic, biomedical and clinical, environmental and behavioral, lifestyle and food preference, and contextual information) along with biospecimens from large cohorts. These invaluable resources offer the potential to advance disease prognosis, risk assessment, and medical and health care through personalized medicine and precision health. However, the vast majority of large studies are conducted in European populations^4,5^, with results that are not optimal for use in other populations. The Taiwan Precision Medicine Initiative (TPMI) is designed to create a Han Chinese cohort to address the needs of almost 20% of the world’s population.

The TPMI, established by Academia Sinica in collaboration with 16 partner medical centers across the country, has built a large cohort of participants who consent to provide DNA samples for genetic profiling and grant access to their electronic medical records (EMR) for studies to develop precision medicine. Of note, EMR access is both retrospective and prospective, ensuring that longitudinal follow-up of each participant is possible. In return, genetic risk profiling results are shared with the participants, with an invitation to participate in follow-up studies to validate disease risk prediction models and risk-based healthcare management guidelines. Key components of the TPMI project include (a) recruitment of a large number of participants from medical facilities in Taiwan; (b) development of population-optimized single nucleotide polymorphism (SNP) arrays; (c) establishment of a dedicated research database for genetic profiles and EMRs; (d) construction of a user-friendly data analysis platform and workplace to facilitate the data retrieval, summary, and visualization for researchers; (e) analysis of genetic profiles and clinical data of the cohort, with a focus on creating algorithms for polygenic risk scores (PRS) to assess common disease risk and pharmacogenetic response; (f) active engagement in public education initiatives aimed at enhancing people’s understanding of genetics and precision medicine.

### Consortium organization

The TPMI is a consortium led by the Steering Committee consisting of the principal investigators of the 17 partner organizations (Academia Sinica and 16 medical centers) that make all project-related decisions by consensus. The Steering Committee is assisted by the Data Access Committee (which evaluates research concepts proposed by consortium members and recommends them for Steering Committee approval prior to data access), the Clinical Application Committee (which establishes best practice health management guidelines based on disease risk), and the Publication Committee (that evaluates abstracts and manuscripts prior to submission for presentation or publication, respectively) (**Fig. S1**). Committee members are drawn from consortium partners and have expertise in clinical research, data analysis, ELSI (Ethical, Legal, and Social Implications), and law. Three teams at the Academia Sinica conduct the study activities: (1) the TPMI Promotion Team coordinates participant recruitment at the partner medical centers, (2) the Genotyping Team performs the genotyping experiments together with 6 hospital-based genotyping teams at consortium hospitals, and (3) the Statistics and Information Technology Team builds the TPMI Data Access Platform (TDAP) and maintains the central database (i.e., TPMI Data Lake) and assist all researchers in data analysis.

### Participant enrollment

Participants were recruited from 16 partner medical centers (encompassing 33 affiliated hospitals) that together serve ∼40% of the population in Taiwan (**Fig. 1**). Informed consent was obtained from the participants while they were enrolled in this study at the hospitals. Participation was offered to all except for those whose peripheral blood cells might harbor non-germline genetic materials: (a) individuals with leukemia who did not achieve remission; (b) individuals who received blood transfusions within the previous six months; (c) individuals who underwent chemotherapy or radiotherapy within the previous 12 months. As of Dec 28, 2023 (TPMI v37 data freeze), 565,390 participants had been enrolled with proper consent.

**Figure 1.**
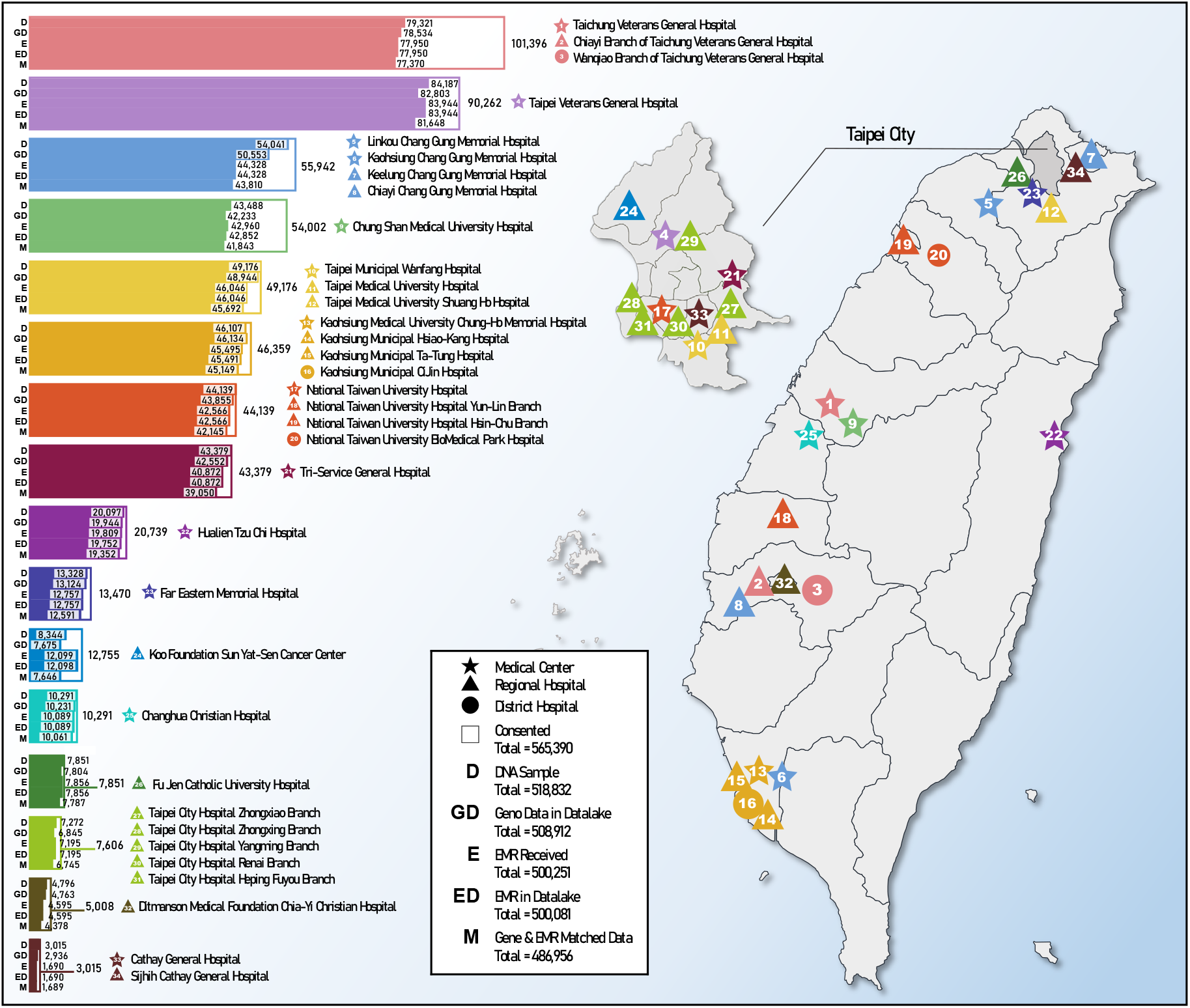
Map of medical centers, their satellite hospitals, and sample sizes.

### Project timeline and milestones

The timeline and milestones are summarized (**Fig. 2**). The TPMI was launched officially in July 2019, when the first participant enrolled with institutional review board (IRB) approval. The TPMI Consortium was established in October 2019, with the Steering Committee constituted at the first consortium meeting in December 2019. Soon after the enrollment reached 200,000 in March 2021, the TDAP was established, and the dataset was made accessible to consortium members in July 2021. The Data Access and Publication Committees were created to facilitate consortium-wide studies and dissemination of results. The Clinical Application Committee was formed in May 2022 to guide the return of results (ROR) to the participants, formulate risk-based healthcare guidelines, and design research studies to validate the results. A SNP array (TPMv1) co-developed with the Taiwan Biobank (TWB) was used initially, and an updated SNP array (TPMv2) was designed and implemented in March 2020. Approximately 3 years after the first enrollment, 500,000 participants were enrolled by September 2022. As of the end of December 2023, there were 565,390 enrolled participants, of which EMRs from 500,081 participants were transferred from the hospital to the TPMI database after the genotyping of 508,912 participants was completed two months before.

**Figure 2.**
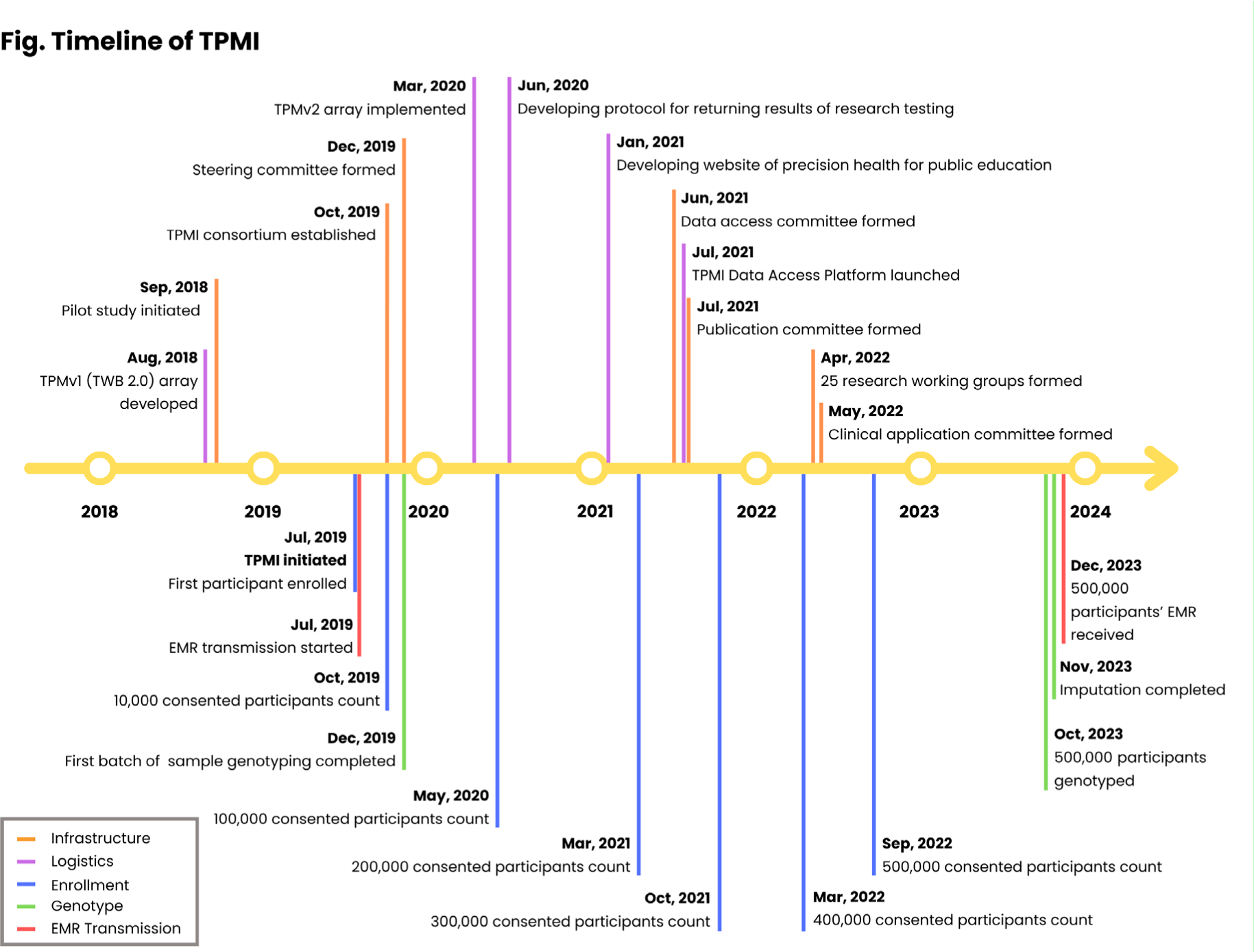
Timeline and milestone of TPMI.

### TPMv1 and TPMv2 custom-designed SNP arrays

In collaboration with the TWB and Thermo Fisher Scientific, TPMI developed the TPMv1 array (686,463 SNPs) specifically for the project. Building on the results from the first year of the project, an updated and optimized TPMv2 array (743,227 SNPs) was subsequently built for the rest of the project. The two TPM arrays were meticulously designed to maximize coverage of the Han Chinese population for genetic studies and to include all previously published disease risk variants. This was done by incorporating (1) the genome-wide imputation grid (GWAS grid) based on the next-generation sequencing (NGS) data from the previous studies^6^, which contained approximately 3,000 samples, including common variants and genetic variants with low minor allele frequencies (MAF) ranging from 1% to 5%; and (2) known disease risk variants from databases such as the GWAS Catalog^7^, ACMG^8^, ClinVar^9^, PharmGKB^10^, and OMIM^11^. After evaluating the performance of the TPMv1 array with data from approximately 100,000 individuals, the TPMv2 array was developed by removing markers with very low MAFs in the Taiwanese population to improve genotyping accuracy on the platform. Additionally, markers used to detect known copy number variations (CNVs) and loss of function (LOF) were added to the array to enhance its utility for genetic analysis (**Table S1**). The TPMv1 and TPMv2 arrays share approximately 495,000 SNPs (**Fig. S2**).

### Genotyping and imputation

Genotyping assays were performed at the National Center for Genome Medicine in the Academia Sinica and 6 partner hospitals (see **Methods–Genotyping experiment and plate normalization**). After quality control measures, we have genotypes for TPMv1 in 99 batches, consisting of 165,596 individuals, and for TPMv2 in 114 batches, comprising 321,360 individuals (486,956 total) with matching EMR data (Version 37). To assess the performance of our phasing and imputation pipeline (see **Methods–Imputation**), 6,000 genotyped variants from 1,000 TPMI individuals on chromosomes 5, 13, and 18 were randomly masked. We assessed imputation quality scores (INFO) and the correlation between imputed and observed genotypes. We found an average correlation of 0.906 for all masked variants, and 96.3% of the masked SNPs have an INFO score >0.7.

### Electronic medical record data

To minimize the burden on the information technology staff of the partner hospitals, the TPMI adopted the strategy of taking EMR data from the hospitals “as is,” except with personal identifying information removed. The TPMI Information Technology Team extracted and standardized the data from diverse hospital data formats into a searchable database to facilitate analysis. For each participant, data from years prior to enrollment, as well as from subsequent hospital and clinic visits, was transmitted to the TPMI database – TPMI Data Lake. 250,000 participants have medical records of five years or more, and 73,000 of them have records of ten years or more. The collected EMR data consist of outpatient records, discharge summaries, laboratory test results, pathology reports, surgery reports, and imaging reports (see **Fig. S3**). Each type of record includes both free-text sections (e.g., Condition Summary in outpatient records; more details in **Table S2**) and predefined structured data (e.g., ICD Diagnosis based on ICD-9 or ICD-10 in outpatient records; more details in **Table S2**). To deal with the many EMR data formats, the Academia Sinica Information Technology Team implemented a series of data quality control (QC) measures during the data import process. These measures include data cleaning, correction, standardization, and extraction. Subsequently, post-QC data was restructured and organized into a custom tabular format, improving search capabilities and overall usability. The team extracted pertinent information from free-text data using natural language processing models or regular expressions for further research analysis. For instance, spaCy models were developed to extract lifestyle data of participants, such as smoking, drinking, and betel nut consumption. At the same time, regular expressions were employed to extract results of cognitive tests, including the Mini-Mental Status Examination, the Cognitive Abilities Screening Instrument, and the Clinical Dementia Rating. In total, 144 EMR variables have been cataloged in the TDAP.

### TPMI Data Analysis Platform (TDAP)

The TPMI genetic and EMR data are encrypted and stored in a centralized, secure research database at the Academia Sinica that is isolated from the internet. The Academia Sinica Information Technology Team developed the TDAP in a protected computing workplace where all data analyses are conducted exclusively at the workplace by the TPMI Consortium members under supervision by the Academia Sinica IT staff and surveillance camera monitoring to ensure that the participants’ data never leave the database. Only summary results from the analyses are disseminated as approved by the TPMI Publication Committee. TDAP enables users to efficiently retrieve data from the TPMI Data Lake for analysis.

Authorized researchers can access TDAP, a secure central database and analysis platform, once their research concepts are approved by the TPMI Data Access Committee and their protocols receive approval from Institutional Review Boards (IRBs). To further support researchers in data exploration, TPMI has developed several platforms. *PheWeb* offers a user-friendly interface for exploring associations between genetic variants and phenotypes, providing access to summary statistics from genome-wide association studies (GWAS) across a wide range of phenotypes and traits. *SNPView* allows users to access information for all loci on the SNP arrays, including the MAF across all TPMI samples, the consistency of loci assessed based on the TWB whole genome sequencing data ^6^, information on genetic variants from ClinVar database ^9^, etc. *DataView* provides information on the number of TPMI participants with specific conditions, laboratory test results, medications prescribed, treatments received, and more.

### Access to data

Access to the TPMI data is based on the consent given by the TPMI participants. First, TPMI researchers registered in TDAP can search metadata and perform initial queries to understand the sample size for specific scientific questions using a system called *PhenoData, which* allows users to input inclusion/exclusion criteria and quickly identify subjects that meet the specified criteria. Researchers can then conduct GWAS and develop polygenic risk scores (PRSs). Upon completion of the analysis, researchers can request to take the summarized results (without any individual information) with them. To date, the TPMI Steering Committee has approved 25 working groups and more than 200 research ideas proposed by TPMI researchers. Researchers outside of the TPMI Consortium with research ideas are encouraged to collaborate with TPMI researchers and pursue studies with the TPMI data.

### Cohort characteristics

Among the 486,956 participants with both genotype and EMR data in the TPMI cohort, there are 217,595 male participants with an average age of 57.4 (standard deviation = 17.5) and 269,361 female participants with an average age of 54.9 (standard deviation = 17.0). The majority of participants fall within the age range of 20 to 90, with over 160 individuals aged over 100 (**Fig. 3A**).

**Figure 3.**
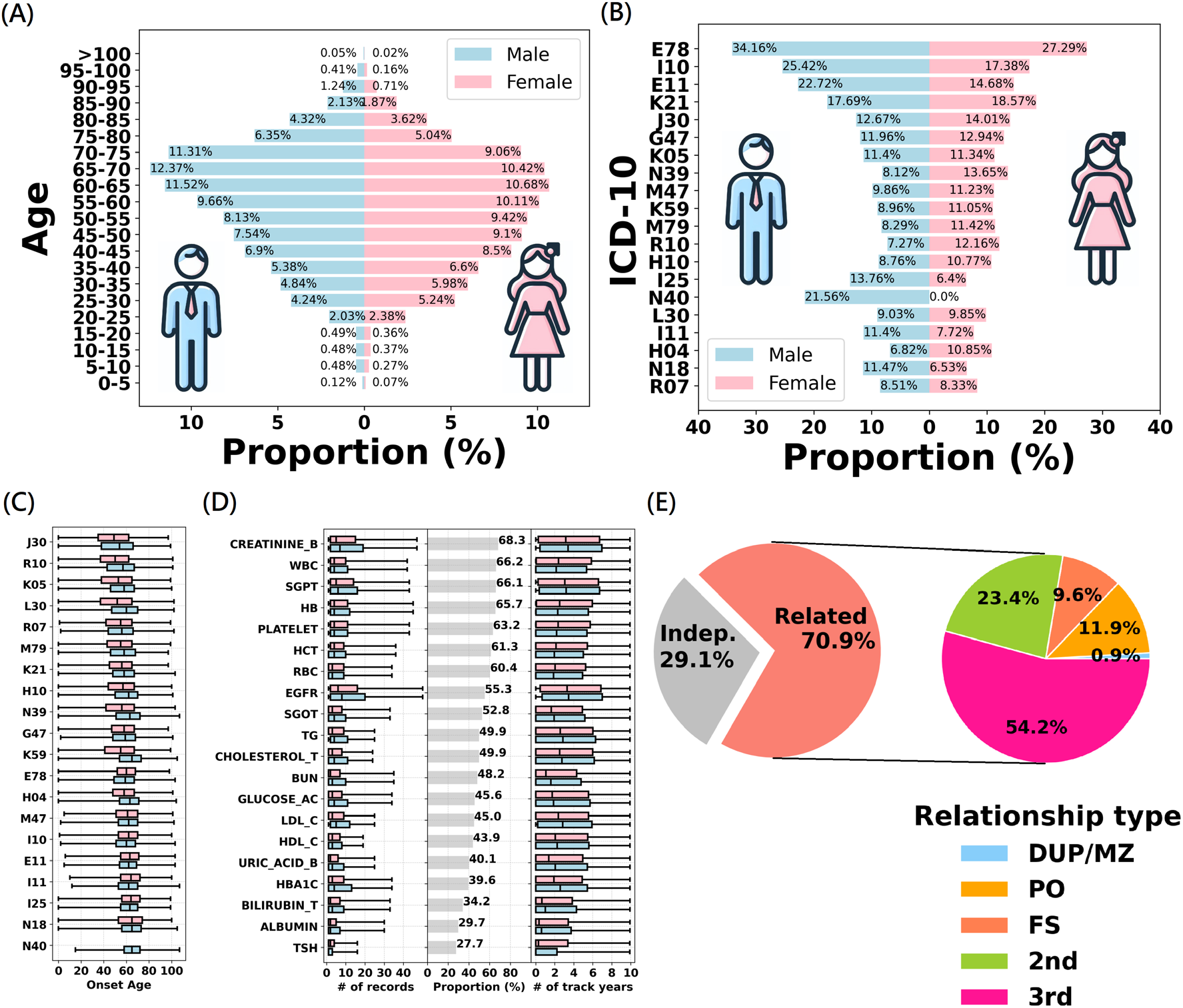
Cohort characteristics. **(A) Sex-specific age distribution. (B) Top 20 most prevalent ICD-10 codes**. This figure displays the proportion of the most frequently observed ICD-10 codes within the study cohort. The list includes codes such as E78 (Disorders of lipoprotein metabolism and other lipidemias), I10 (Essential hypertension), E11 (Type 2 diabetes mellitus), K21 (Gastro-esophageal reflux disease), J30 (Vasomotor and allergic rhinitis), G47 (Sleep disorders), K05 (Gingivitis and periodontal diseases), N39 (Other disorders of the urinary system), M47 (Spondylosis), K59 (Other functional intestinal disorders), M79 (Other and unspecified soft tissue disorders), R10 (Abdominal and pelvic pain), H10 (Conjunctivitis), I25 (Chronic ischemic heart disease), N40 (Enlarged prostate), L30 (Other and unspecified dermatitis), I11 (Hypertensive heart disease), H04 (Disorders of the lacrimal system), N18 (Chronic kidney disease), and R07 (Pain in throat and chest). **(C) Onset age of the top 20 most prevalent diseases**. This figure displays the sex-specific onset age of the disease. For each disease, the onset ages of males (in blue) and females (in pink) are presented in a box-whisker plot. The diseases are arranged based on the median onset age. **(D) Top 20 most prevalent lab tests**. This figure provides a comprehensive overview of data availability across 20 laboratory tests, including CREATININE_B (Creatinine of blood), WBC (White Blood Cell), SGPT (S-GPT/ALT), HB (Hemoglobin), PLATELET (Platelet Count), HCT (Hematocrite), RBC (Red Blood Cell), EGFR (Estimated Glomerular Filtration Rate), SGOT (S-GOT/AST), TG (Triglyceride), CHOLESTEROL_T (Total Cholesterol), BUN (Blood Urea Nitrogen), GLUCOSE_AC (Glucose(AC)), LDL_C (Low-Density Lipoprotein Cholesterol), HDL_C (High-Density Lipoprotein Cholesterol), URIC_ACID_B (Uric Acid (Blood)), HBA1C (Hemoglobin A1c), BILIRUBIN_T (Bilirubin (Total Value)), ALBUMIN (Albumin Value), and TSH (Thyroid-Stimulating Hormone (EIA/LIA) Value. The three panels from the left-hand side to the right-hand side indicate the sex-specific distribution of the number of records for each individual, the proportions of samples with available lab test data, and the distribution of the average follow-up years of the lab tests. Note that the number of records is winsorized, i.e., values exceeding the 95th percentile were replaced by the 95th percentile value. **(E) Familial relativeness**. The first pie chart displays the proportions of related and unrelated samples. The second pie chart further illustrates the proportions of inferred relatedness, including duplicate (DUP) or monozygotic twin (MZ), parent-offspring (PO), full-sibling (FS), second-degree (2nd), and third-degree (3rd) relativeness.

Among the participants with ICD-10 codes, the top five prevalent diseases include disorders of lipoprotein metabolism and other lipidemias (E78, 30.3%), essential hypertension (EHT) (I10, 21.0%), type 2 diabetes (T2D) (E11, 18.3%), gastroesophageal reflux disease (K21, 18.2%), and vasomotor and allergic rhinitis (J30, 13.4%) (**Fig. 3B**).

Consequently, metabolic syndrome emerges as the most prevalent health conditions. Noteworthy sex differences in the diagnosis proportion were observed among the top 20 diseases, with exceptions noted in gingivitis and periodontal diseases (K05) (p = 0.56). The data suggest that age- and sex-adjustment is necessary in subsequent genetic association analyses. These top 20 prevalent diseases have differential onset ages, where vasomotor and allergic rhinitis (J30) has the youngest average onset age of 49.4 (standard deviation = 18.4) and enlarged prostate (N40) has the oldest average onset age of 65.5 (standard deviation = 11.1) (**Fig. 3C**).

Among 486,956 participants with laboratory test results of urine and blood samples, 68.3% have creatinine test records, averaging 5 test records per person and demonstrating an average follow-up duration of 3 years (**Fig. 3D**). The subsequent 2^nd^ to 5^th^ frequent lab tests include white blood cell (66.2%), serum glutamic pyruvic transaminase (66.1%), hemoglobin (65.7%), and platelet count (63.2%) (left panel in **Fig. 3D**). These prevalent tests are integral components of standard diagnostic and preventive care panels, and their data availability may also be linked to the prevalent health issues and chronic conditions. For instance, creatinine levels in the blood are commonly utilized to assess kidney function, associated with prevalent conditions in Taiwan, such as urinary tract issues, hypertension, or diabetes.

Familial relatedness analysis using kinship coefficient and identity by descent (see **Methods–Familial relativeness analysis**) reveals that 70.9% of participants can identify their third-degree or closer relatives among other TPMI participants, with the distribution of different levels of relativeness provided (**Fig. 3E**). For genetic association analysis, which assumes sample independence, it is necessary to select unrelated representative samples from each family. However, this approach results in a substantial decrease in sample size.

Alternatively, employing a generalized mixed effect approach, which analyzes sample correlation by considering a random effect – such as SAIGE ^12^ for case-control studies and BOLT-LMM ^13^ for quantitative trait studies – can be used for genome-wide association study (GWAS) without a loss in sample size.

### Population structure

The population structure of the TPMI cohort was analyzed against external resources with known population information, including TWB^14,15^, Simons Genome Diversity Project (SGDP)^16^, and the 1000 Genomes Project (1KG)^17^ (see **Methods–Population structure analysis**). Given the major influx of people from mainland China to Taiwan around 1950^18,19^, a separate principal component analysis (PCA) was conducted specifically for TPMI participants born before 1950, referred to as “<1950,” with a sample size of n = 70,708 (14.6%). The first two principal components (PCs) were used to construct a reference coordinate system, and subsequently, all other participants were projected onto the reference coordinate system.

TPMI participants born after 1950 exhibit a more diverse distribution, reflecting the historical intermarriage and genetic admixture between the major ethnic groups (the Minnan, Hakka, and the Mainlanders) and minor indigenous groups (**Fig. 4**). It is worth noting that TPMI and TWB (another large cohort project in Taiwan with a community-based design) have similar patterns in the PCA plot because the majority of both cohorts are of Han Chinese ancestry.

**Figure 4.**
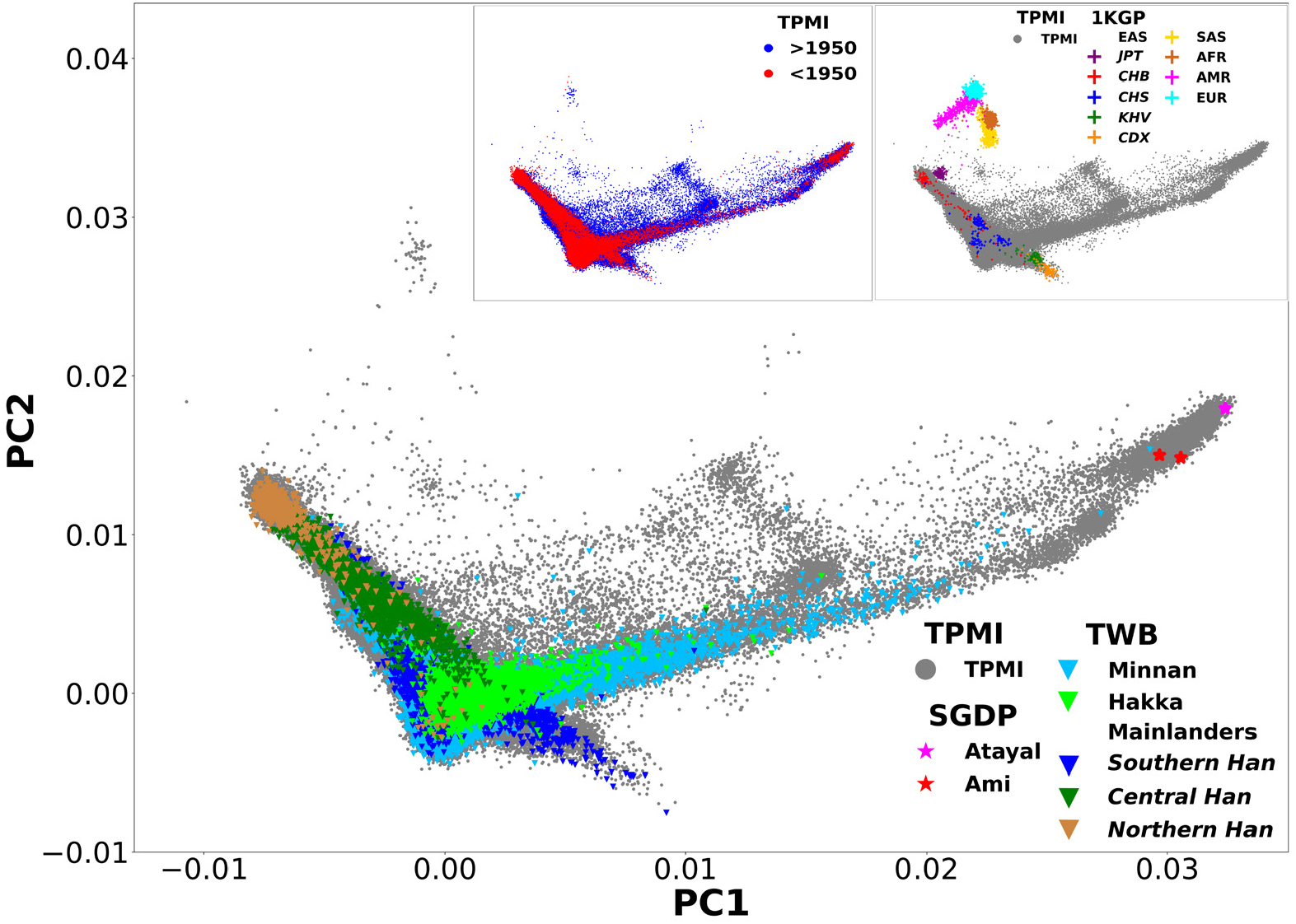
Population structure. The population structure of the TPMI cohort was assessed against samples from the Taiwan Biobank (TWB), the Simons Genome Diversity Project (SGDP), and the 1000 Genomes Project (1KG) using principal component analysis. The top left subfigure presents a contrast of the TPMI participants born before and after 1950. The top right subfigure presents a contrast between the TPMI and 1KG datasets. The main figure exhibits a comparison of the TPMI cohort with the Taiwan Biobank (TWB) and two Taiwan indigenous tribes included in the Simons Genome Diversity Project (SGDP).

The first PC primarily distinguishes between the Han Chinese and the indigenous groups. This is confirmed by comparing the TPMI data against individuals from two indigenous tribes (Ami and Atayal) in the SGDP dataset (**Fig. 4**). Ami and Atayal individuals can be further separated by higher-order PCs.

The second PC is correlated with latitude, where higher PC2 scores corresponded to northern latitudes, and lower scores indicated southern latitudes. This pattern is observed in the TWB dataset (Southern, Central, and Northern Han from mainland China) and also in the 1KG project dataset (East Asians (EAS)= CDX, KHV, CHS, CHB, and JPT). PCA and population admixture analysis allow us to adjust for population structure when conducting genetic studies and exclude non-Han Chinese individuals from subsequent GWASs.

Homozygosity analysis (refer to **Methods–Homozygosity analysis**) reveals that individuals closer to the indigenous and non-East Asian ethnic groups exhibit higher homozygosity (**Fig. S4A**). The main explanation is that the SNPs on the custom-designed Han Chinese TPM arrays are more often monomorphic in indigenous or non-Han Chinese groups, also supported by the analysis result based on the SNPs on the TPM arrays showing that cohorts with an East-Asian ancestry (TPMI, TWB, and EAS) exhibited a lower homozygosity level compared to other non-East-Asian ethnic groups (**Fig. S4B**). Note that the homozygosity distributions differ from those derived from whole-genome sequencing analysis in other ethnic groups^20^, serving as a reminder that direct use of the SNP panels on the TPM arrays is well-suited to the genetic research of the East-Asian population but may not be as useful for non-East-Asian populations. On average, in the TPMI cohort, the median homozygosity rate was 0.801 (standard deviation = 0.003), with 1,197 individuals exceeding six times the interquartile range.

### GWAS, QTL mapping, and sample size evaluation

As to sample size evaluation (**Methods–Sample size evaluation**), to detect a SNP that has odds ratio from 1.1 – 2.0 for a condition based on MAF of 0.01 – 0.25, case/control ratio of 1:4, and significance level of 5×10^−8^, the sample size required for attaining a statistical power of 0.8 was calculated by Quanto ^21^ (**Fig. 5A**). To detect a quantitative trait locus (QTL) that has a beta coefficient from 0.02 – 0.20 for a condition based on MAF of 0.1 – 0.25, and significance level of 5×10^−8^, the sample size required for attaining a statistical power of 0.8 was calculated by Quanto (**Fig. 5B**).

**Figure 5.**
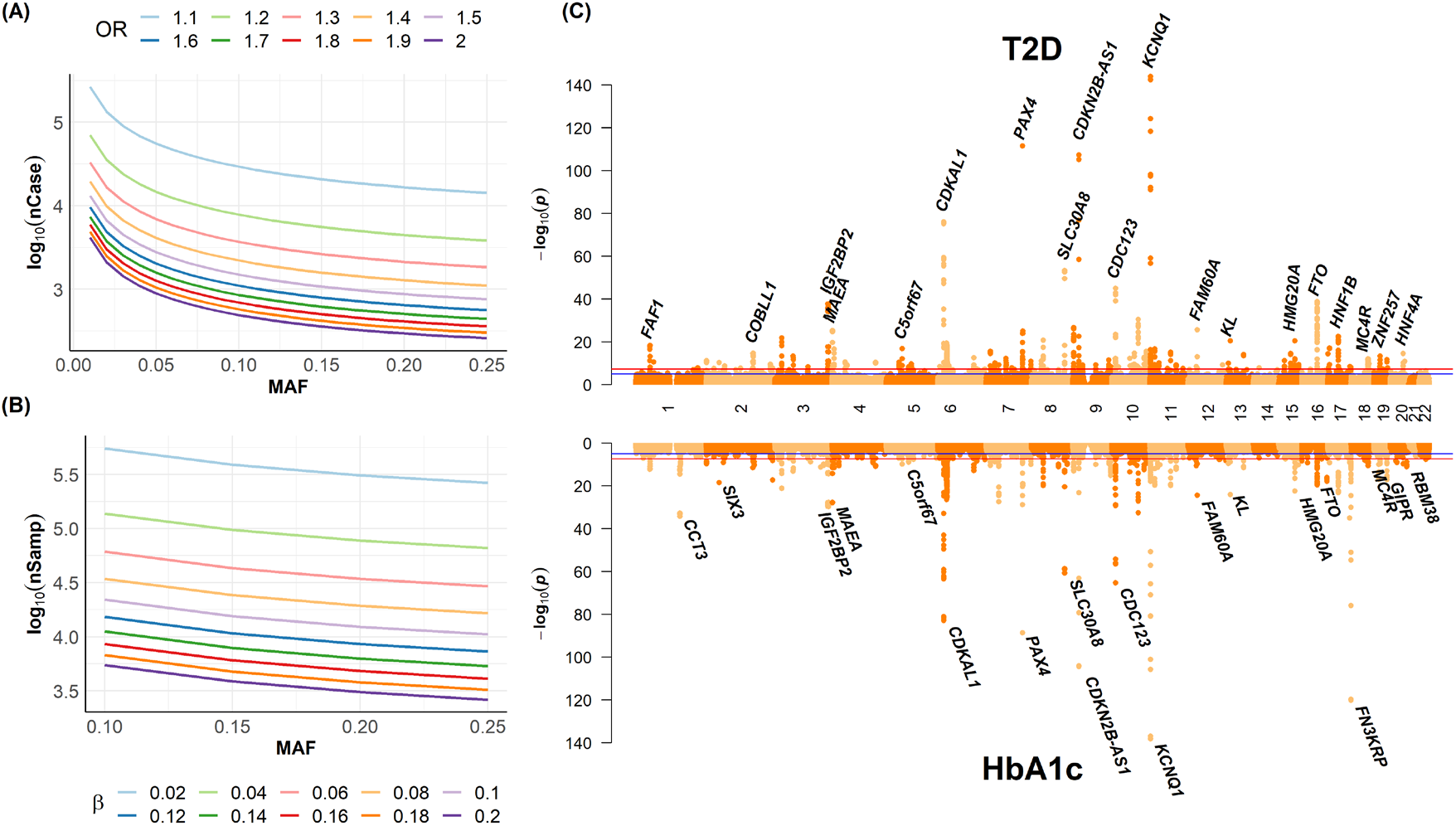
Sample size evaluation and examples for GWAS and QTL mappings. **(A) Sample size calculation for a case-control study**. The horizontal axis is minor allele frequency (MAF). The vertical axis is the number of cases on a scale of log 10. Curves with different colors reflect different effect sizes. **(B) Sample size calculation for quantitative trait locus (QTL) study**. The horizontal axis is minor allele frequency (MAF). The vertical axis is the number of participants on a scale of log 10. Curves with different colors reflect different effect sizes (i.e., beta values). **(C) Miami plot of the GWAS for Type 2 Diabetes (T2D) and QTL mapping for HbA1c**. The red (blue) reference line indicates a significance level of *p* = 5×10^−8^ (*p* = 1×10^−5^). SNPs with *p* < 1×10^−8^ were identified, and the names of the genes harboring these significant SNPs are displayed.

For example, the prevalence of diabetes mellitus in Taiwan from 2017-2020 was 11.3%.^22^ The TPMI cohort has 72,659 T2D cases, sufficient to detect T2D-associated SNPs with an odds ratio higher than 1.1 and MAF higher than 0.05. After rigorous data quality control (refer to **Methods–Quality control** and **Figs. S5 – S6**), a preliminary GWAS of T2D (refer to **Methods–Genome-wide association study**) in the TPMI cohort replicate previous findings such as *Potassium voltage-gated channel subfamily Q member 1* (*KCNQ1*) (*p* = 9.78×10^−145^), *Diabetes-Linked Transcription Factor paired box 4* (*PAX4*) (*p* = 2.67×10^−112^), and *CDKN2B antisense RNA 1* (*CDKN2B-AS1*) (*p* = 5.14×10^−108^) for T2D (n = 52,290 cases and 192,817 controls, top panel in **Fig. 5C**).

In addition, regarding quantitative trait locus (QTL) analysis, the TPMI cohort has 192,701 participants who have HbA1c records, sufficient to detect HbA1c-associated QTLs with a beta coefficient higher than 1.1 and MAF higher than 0.05. Preliminary GWAS of HbA1c identified the same genes in the GWAS for T2D, such as *KCNQ1* (*p* = 6.77×10^−139^), *CDKN2B-AS1* (*p* = 5.08×10^−105^), and *PAX4* (*p* = 3.18×10^−84^), and the additional genes, such as *fructosamine 3 kinase related protein* (*FN3KRP*) (*p* = 9.11×10^−121^) and *chaperonin containing TCP1 subunit 3* (*CCT3*) (*p* = 7.46×10^−35^) (n = 140,259, bottom panel in **Fig. 5C**).

In another example, GWAS for essential hypertension (EHT; n = 71,548 cases and 130,561 controls, **Fig. S7A**) and QTL mappings for systolic blood pressure (SBP; n = 241,667, **Fig. S7B**) and diastolic blood pressure (DBP; n = 241,646, **Fig. S7C**) simultaneously identified *Fibroblast Growth Factor 5* (*FGF5*) (*p* = 2.41×10^−69^), *Fat Mass and Obesity-Associated* (*FTO*) (*p* = 8.74×10^−19^), and *RAL Guanine Nucleotide Dissociation Stimulator Like 3* (*RGL3*) (*p* = 4.49×10^−13^), along with other genes specific to each of the three GWAS and QTL mappings. Additional results from GWASs and phenome-wide association studies (PheWASs) for important diseases in Taiwan and their subtypes, as well as quantitative traits, are reported (Chen et al., Population-Specific Polygenic Risk Scores Developed for the Han Chinese, Submitted to MedRxiv, DOI pending). Generalized mixed-effect analysis using SAIGE^12^ also exhibits similar results (**Fig. S8**). These results elucidate the ability to uncover the genetic underpinnings of complex disorders and traits such as T2D and EHT using the TPMI cohort.

### Polygenic risk score

In the example of T2D, two multi-ancestry PRSs were constructed (refer to **Methods**– **Polygenic risk score**). The first method applied the summary genetic effects from meta-GWASs that comprised one million participants with East Asian, European, and South Asian ancestry in the DIAGRAM Consortium^23^ achieved an area under the receiver operating characteristic curve (AUC) of 0.65 in both the training and the testing datasets. After incorporating age and sex, the AUC is further increased to 0.83 in both the training and testing datasets (**Fig. 6A**). The second PRS based on the effect sizes in PGS (PGS002308) ^24^ achieved an AUC of 0.65 in the training and testing datasets, and after incorporating age and sex, the AUC is further increased to 0.83 in both training and testing datasets (**Fig. S9A**). The positive dose-response correlation between the PRS level and T2D odds ratio (**Fig. 6B** and **Fig. S9B**) and enhanced PRS after incorporation of demographic factors such as age and sex (**Fig. 6A** and **Fig. S9A**), highlighting the potential of PRS in identifying individuals at heightened risk for T2D, thus enabling targeted interventions and more precise therapeutic strategies. Additional results regarding PRS for important diseases in Taiwan and their subtypes are reported (Chen et al., Population-Specific Polygenic Risk Scores Developed for the Han Chinese, Submitted to MedRxiv, DOI pending).

**Figure 6.**
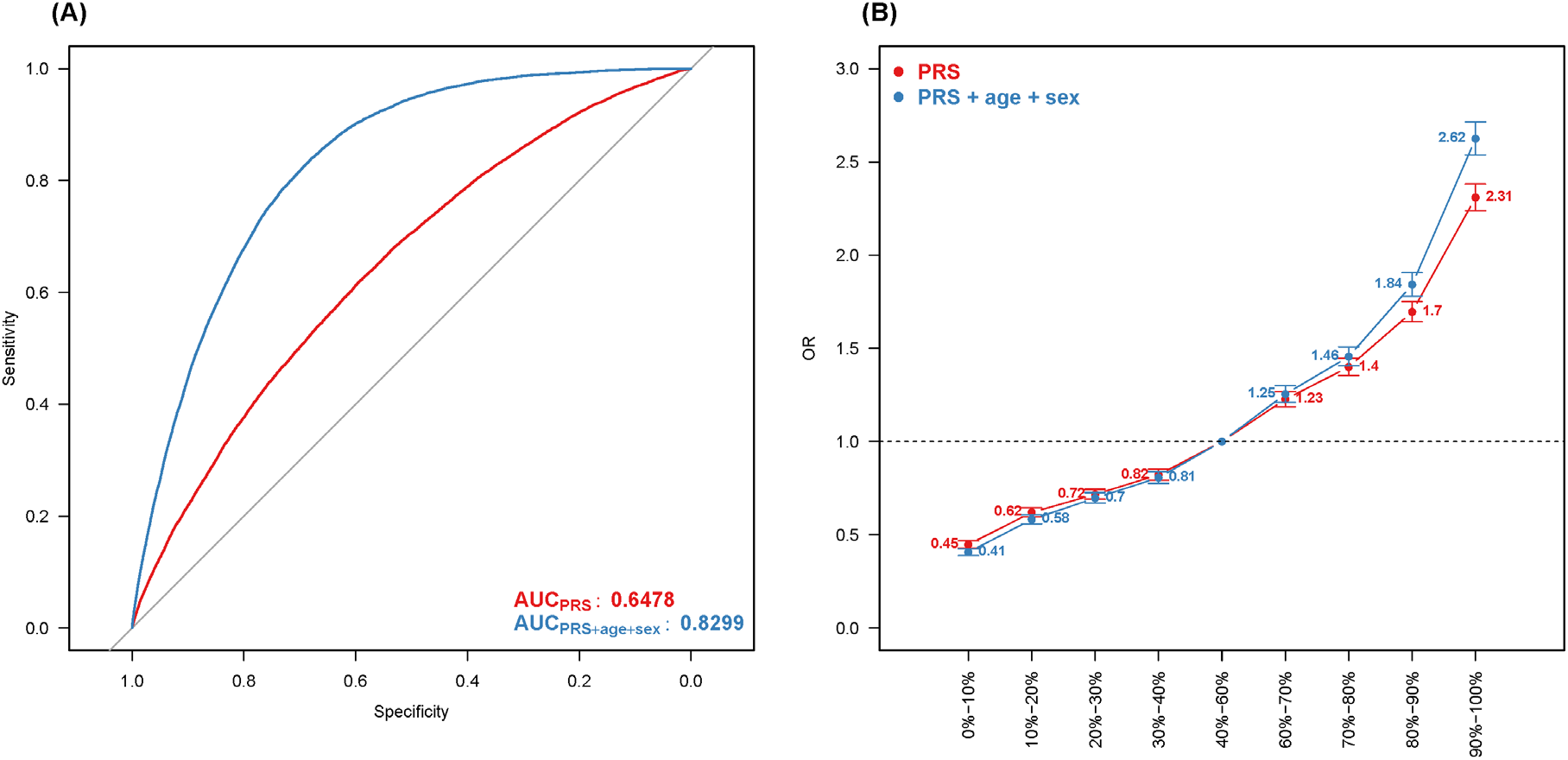
Polygenic risk score analysis. **(A) Area under the receiver operating characteristic curve (AUC)**. AUC of PRS (red curve) and AUC of PRS, age, and sex (blue curve) are displayed. **(B) Dose-response effect of PRS levels on the odds ratio of T2D**. Dose-response effect of PRS (red line) and dose-response effect of a combination of PRS, age, and sex (blue curve).

## Discussion

The TPMI has reached a cohort size of >500,000 participants of Han Chinese ancestry, where genetic and EMR data are available for analysis. As such, it is the largest non-European cohort of its kind. The genetic homogeneity and richness of clinical data (from years prior to enrollment plus those from future hospital visits) make the cohort highly valuable for genetic and epidemiological research. As reported elsewhere, results from large-scale GWASs, common disease risk prediction (based on PRS) (Chen et al., Population-Specific Polygenic Risk Scores Developed for the Han Chinese, Submitted to MedRxiv, DOI pending), deleterious variants (Huang et al., Deleterious Variants Contribute Minimal Excess Risk in Large-Scale Testing, Submitted to BioRxiv, DOI pending), and pharmacogenetics (Wei et al., Clinical Impact of Pharmacogenetic Risk Variants in a Large Chinese Cohort, Submitted to MedRxiv, DOI pending) in the TPMI cohort have real-world implications. The commitment to return study results to the participants and to involve them in future clinical research will facilitate the validation of precision medicine approaches in healthcare management.

However, the TPMI cohort has two sets of limitations. Firstly, the quantity and quality of the clinical data are not perfect. While the TPMI participants grant us access to “all” their clinical data, the project lacks the resources and manpower to retrieve hospital data in archival storage, limiting access to entire data before their enrollment in the project. In addition, because many patients receive care from multiple hospitals and clinics under Taiwan’s National Health Insurance Program, clinical data from sources outside the hospital through which a participant enters the project are unavailable to us. These circumstances result in incomplete clinical data, leading to cases where the age of disease onset, test results, treatments prescribed, and drug responses are missing. Secondly, due to technical constraints, the genetic risk profile data do not encompass some known risk variants.

Although many known risk variants are included in the SNP array, some cannot be genotyped because suitable probes cannot be designed on the array. Furthermore, the genotyping accuracy of SNP arrays is low when the MAF of the marker is less than 0.1%, making it challenging to confidently type a substantial fraction of the known risk variants in the TPMI cohort, as they are scarce.

### Future directions

The primary focus of the TPMI is to develop algorithms to predict disease risk for as many conditions as the cohort can support. Once developed, the algorithms must be validated in real-world settings before being adopted for the population. The TPMI cohort will be very useful in this regard. For common diseases that affect older individuals, there will be some in the TPMI cohort who are in the high-risk group for each disease but are yet unaffected. Following these individuals, especially those approaching the expected age of onset, can provide a measure of the predictive power of the algorithms. Furthermore, for some high-risk groups where health management strategies are available for the diseases in question, a trial comparing those who follow the risk-lowering guidelines versus those under the standard of care will determine if genetic risk-guided health management is beneficial. For example, those with high cancer risk can be enrolled in an early screening program, and those with high stroke risk can be enrolled in a stroke prevention program that includes blood pressure control and smoking cessation. To make the cohort even more helpful, additional resources will be sought to retrieve archival clinical data from the hospitals through which the participants join the TPMI and obtain consent from the participants to extract data from the National Health Insurance Database^25,26^ and other hospitals and clinics where they receive care. With this enhanced dataset and longitudinal follow-up data, the TPMI cohort can be studied for years to come.

In addition to TPMI, the TWB^14,15^ and the China Medical University Hospital (CMUH)^27^ represent two additional large cohorts for genetic studies in Taiwan. TWB and CMUH have recruited two hundred thousand and one hundred seventy thousand participants, respectively. TWB aimed for broad recruitment across Taiwan, while CMUH focused more regionally. The integration of TPMI, TWB, and CMUH data forms one of the largest cohorts for genetic studies globally, significantly enhancing statistical power for GWAS and PRS for the East Asian population. However, this integration also presents challenges in genetic analysis due to differences in study design and data collection among the three large cohorts. For instance, TWB collected self-reported disease records through questionnaires rather than through medical diagnosis in EMR, whereas CMUH primarily cares for patients from central Taiwan. These differences among the three cohorts present challenges in data analysis. A meta-analysis based on summary statistics may offer a viable approach, but careful adjustment for potential confounders and background differences among the three cohorts is necessary. Advanced methods should be developed.

RoR is crucial to empower participants through engagement and education, raise public awareness of precision health, and facilitate the establishment of infrastructure for clinical implementation. For the TPMI project, an RoR platform for 83 genetic conditions has been developed with custom-designed RoR webpages by each hospital. The content includes disease-related variants and pharmacogenetics-related variants. For disease-related variants, founder mutations, or pathogenic variants with multiple evidence (following The American College of Medical Genetics and Genomics (ACMG) and National Comprehensive Cancer Network (NCCN) guidelines) related to cancer, dermatology, endocrine, hearing loss, hematology, metabolism, neurology, and ophthalmology are interpreted (**Table S3**). For pharmacogenetics-related variants, actionable variants from US FDA and Clinical Pharmacogenetics Implementation Consortium (CPIC) are selected to report, and the therapeutic ranges of involved drugs include anesthesiology, anti-inflammatory, cardiology, endocrinology, gastroenterology, hematology, hyperuricemia, infectious diseases, muscle tenderness, neurology, oncology, psychiatry, toxicology, transplantation, etc. (**Table S3**).

Genetic consultation is essential in guiding individuals, families, and society through life-changing decisions based on genetic information, especially in implementing ROR. Prioritizing genetic consultation within the TPMI will address several vital considerations, including facilitating informed decision-making, safeguarding the privacy and confidentiality of genetic data, navigating ethical complexities, providing psychosocial support for emotional challenges, and ensuring equitable access to precision health services across diverse demographic populations. The TPMI is positioned to transform healthcare by blending scientific progress with ethical awareness, particularly through advancing precision health. Recognizing the crucial role of genetic consultation in managing ELSI in Taiwan, the TPMI will be dedicated to harnessing the power of genetic data while valuing the guidance provided by genetic counselors in navigating ELSI complexities.

## Methods

### Genotyping experiment and plate normalization

Genomic DNA was purified automatically from 200 μl of whole blood with QIAsymphony DSP DNA Mini Kit (QIAGEN, Germantown, MD). 15 μl of genomic DNA at 50 ng/μl was subjected to genotyping using Axiom TPMv1 (Axiom TPM) or TPMv2 Array (Axiom TPM2) (Thermo Fisher Scientific, Waltham, MA) according to the manufacturer’s instruction. Genotyping assays were performed at the National Center for Genome Medicine in the Academia Sinica, Taipei, Taiwan (https://ncgm.sinica.edu.tw/) and 6 partner hospitals, including Center of Applied Genomics at Kaohsiung Medical University, Kaohsiung, Taiwan; Precision Medicine Center at Taichung Veterans General Hospital, Taichung, Taiwan; Chang Gung Memorial Hospital, New Taipei City, Taiwan; Chuanghua Christian Hospital, Chuanghua, Taiwan; Hualien Tzu Chi Hospital, Hualien, Taiwan; and Taipei Medical University, Taipei, Taiwan. The genotype calling was performed for approximately 3,000 individuals (ranging from 2,304 to 3,936) per batch using Applied Biosystems™ Array Power Tools (APT) as part of the Best Practices Workflow in National Center for Genome Medicine at Academia Sinica. Each batch included arrays from consecutive assays at individual centers to minimize the potential batch effect. Individuals with a call rate ≤98% were genotyped again to improve the call rate.

We also performed plate normalization to examine whether there were differences in the signal distribution of 96-well plates within the same calling batch for each marker.

Variations in signal intensity across different plates can lead to misjudgment by the clustering algorithm during the genotype calling stage. The normalization procedure helps mitigate such issues, reduce the impact of signal intensity variations, and improve clustering accuracy. We calculated the allele frequency of each marker to identify any abnormalities and, based on these calculations, determined whether normalization was required.

### Imputation

Whole genome sequencing data using Illumina HiSeq and Novaseq from 1,498 subjects from the TWB were used as imputation references ^6^. The sequencing reads were aligned to the human genome reference GRCh38 using BWA ^28^. Variants were called jointly with DeepVariant ^29^. Read-based phasing with WhatsHap ^30^ initially, followed by population-level phasing with SHAPEIT4 for better accuracy ^31^. Removal of variants with minor allele count <2, Hardy-Weinberg equilibrium test p-value <1×10^−10^, or missing rate >5% resulted in 22.44 million genetic variants in the imputation reference panel. SHAPEIT5 and IMPUTE5 were applied to all genotyped individuals for haplotype phasing and genome imputation ^32,33^.

### Familial relativeness analysis

For the samples that passed sex, inconsistent duplicated EMR, and call-rate check (**Fig. S3**), we utilized a dataset comprising 485,925 individuals and 68,741 unlinked SNPs to estimate familial relationships using KING software (version 2.2.7)^34^. These SNPs were selected based on criteria where the MAF was greater than 5%, the SNP call rate was at least 99%, and pairwise linkage disequilibrium was less than 0.3 within a sliding window of 5 Mb. The inference of close relationships, such as duplicates/monozygotic twins (Dup/MZ), parent-offspring (PO), full siblings (FS), second-degree (2nd), and third-degree (3rd), using the ‘-related’ option, which estimates kinship coefficient by the proportion of genomes shared Identical-by-descent (IBD).

### Population structure analysis

The population structure of the TPMI cohort was assessed against external resources with known population information from various genetic projects, including the TWB ^14,15^, the Simons Genome Diversity Project (SGDP) ^16^, and the 1000 Genomes Project (1KG) ^17^. The TWB dataset encompassed 83,664 individuals, consisting of 68,023 with Minnan ancestry, 11,549 with Hakka ancestry, and 4,092 Han Mainlanders, further categorized into 1,681 Southern Han, 1,606 Central Han, and 805 Northern Han based on self-reported birth geographic regions. The SGDP dataset included 3 individuals from two Taiwan indigenous tribes, namely 1 Atayal people and 2 Ami people, to assess the genetic contribution of indigenous populations in Taiwan. Within the 1KG dataset, there were 3,202 individuals representing 26 global populations across five continents – Africa (AFR), Americas (AMR), East-Asia (EAS), South-Asia (SAS), and Europe (EUR). This dataset comprised 893 with AFR ancestry, 490 with AMR ancestry, 585 with EAS ancestry, 601 with SAS ancestry, and 633 with EUR ancestry. The EAS-ancestry group consisted of 104 Japanese in Tokyo, Japan (JPT), 103 Han Chinese in Beijing, China (CHB), 163 Southern Han Chinese, China (CHS), 93 Chinese Dai in Xishuangbanna, China (CDX), and 122 Kinh in Ho Chi Minh City, Vietnam (KHV).

Principal component analysis (PCA) was conducted based on a set of 234,255 autosomal SNPs common to both TPMv1 and TPMv2 SNP arrays. These SNPs passed stringent quality control measures, including an MAF greater than 1% and a call rate exceeding 99% (**Fig. S3**). Additionally, SNPs with an inter-marker linkage disequilibrium of r2 less than 0.2 were chosen. The analysis comprised 70,708 TPMI individuals who passed stringent sample quality control criteria (**Fig. S3**) and were born before 1950, referred to as “<1950.” The first two principal components (PCs), which explained 43.9% and 18.8% of the genetic variation, respectively, were derived from the analysis involving a total of 10 PCs, as calculated from their corresponding eigenvalues. These components were utilized to construct a reference coordinate system. Subsequently, all other participants, including the individuals of “>1950” in TPMI, TWB, and SGDP, were projected onto this reference coordinate system. For computational efficiency, PCA and the top 10 PCs were generated using the fastPCA version (--pca approx) in PLINK 2.0.

### Homozygosity analysis

The homozygosity rate of each individual was calculated by using PLINK 2.0 (--het) based on 479,610 autosomal SNPs shared in TPMv1 and TPMv2. In order to observe a homozygosity pattern in the TPMI, a smoothing homozygosity rate was calculated as follows. PCA based on genotype data (**Fig. S4A**) was performed, and the coordination of the first two PCs was divided into 150 evenly-spaced partitions. The partitions contained no individuals were removed, resulting in 4,739 partitions remaining. In each PC partition, homozygosity rates of individuals were calculated and visualized in a heat map (**Fig. S4A**). In addition, cross-ancestry distributions of individual’s whole-genome homozygosity rates were visualized in violin plots for comparison (**Fig. S4B**).

### Sample size evaluation

We used QUANTO 1.2.4 (Quantitative Trait Loci ANalysis TOol)^21^ to calculate the required sample sizes for the identification of a disease-associated SNP in a GWAS or QTL in a QTL mapping to attain a statistical power of 0.8 under a genome-wide significance level of 5×10^−8^ for various situations. For binary traits, based on a logistic regression, we considered SNPs with an MAF ranging from 0.01 to 0.25 and an odds ratio ranging from 1.1 to 2.0, under the ratio of cases to controls of 1-to-4. For quantitative traits, based on a linear regression, we considered SNPs with an MAF ranging from 0.01 to 0.25 and an effect size ranging from 0.02 to 0.2.

### Quality control

We performed sample and SNP quality control using PLINK in cooperation with KING and R^35^ (refer to **Fig. S5**). Initially, there were 486,956 participants with both EMR and genotyping data in either TPMv1 array (n = 165,956) or TPMv2 array (n = 321,360), and 479,610 shared autosomal SNPs genotyped on both arrays. We began by excluding SNPs in specific batches of participants with significantly different allele frequencies compared to other batches.

Subsequently, we sequentially identified and removed problematic participants: 705 with inconsistencies between EMR-recorded gender and genetic gender determined by homozygosity pattern of the X chromosome; 307 assembled into 2 to 3-person groups by highly similar genomes had inconsistent EMR records for either genders or birth dates; 19 with a low genotyping call rate (GCR) of <0.95 (--mind 0.05); 8,123 were with excessive or reduced autosomal heterozygosity rate (--het) more than three standard deviations away from the mean of heterozygosity rates; 111,489 with equal to or higher than 2nd-degree cryptic relations with other participants estimated by KING; 1,135 deviated from 99.99% confidence bands (R {car}) of the first two principal components of genetic relationship matrix (GRM) projected onto the 1000 Genomes project dataset (--score). Here, we retained 485,925 participants who passed at least the call-rate check for the population structure investigation and 476,449 related and 365,178 independent participants for the subsequent GWAS and PRS analyses. Based on unrelated participants, we hierarchically excluded SNPs for each studied trait by GCR of <0.95 (--geno 0.05) or those that failed the nonrandom missingness test for a binary trait (--test-missing), minor allele frequency (MAF) of <0.01 (-- maf 0.01), and p-value of the Hardy-Weinberg equilibrium test at the Bonferroni’s level (-- hwe). Finally, approximately 440,000 filtered SNPs remained for each trait.

### Genome-wide association study

We conducted genome-wide association studies (GWASs) for two binary disease traits: type 2 diabetes (T2D) and essential hypertension (EHT). Additionally, we performed quantitative trait locus (QTL) mappings for three quantitative traits: glycated hemoglobin (HbA1c), systolic blood pressure (SBP), and diastolic blood pressure (DBP).

For binary disease traits, we defined disease status using ICD-10 codes from EMR records with lab tests. A T2D patient was defined as having at least 10 records of ICD10-code E11 or had HbA1c ≥6.5% and fasting glucose (FS) ≥126 mg/DL. A non-T2D control was defined as having none of the ICD-10 codes related to diabetes mellitus (DM) and none of the records with HbA1c over 5.6% or FS over 100 mg/DL. Similarly, an EHT patient was defined by the ICD10-code I10, systolic blood pressure (SBP) ≥120 mmHg, or diastolic blood pressure (DBP) ≥80 mmHg. A non-EHT control did not meet any of the aforementioned EHT inclusion criteria.

Firth logistic regression with age, sex, and 10 principal components (PCs) for ancestry adjustments was implemented by PLINK 2.0 (--glm) on an independent sample dataset. In addition, a logistic mixed-effect model adjusted by the same covariates (i.e., age, sex, and 10 PCs) was implemented by SAIGE 1.1.9 on a related-samples dataset. For quantitative traits, we first applied the inverse normal transformation^36^ to the residuals obtained by regressing the quantitative trait against the aforementioned covariates. Subsequently, a linear regression was implemented by PLINK on an independent-samples dataset, and a linear mixed-effect model was implemented by SAIGE on a related-samples dataset.

### Polygenic risk score

We computed multi-ancestry PRS for T2D using a Bayesian approach, PRS-CSx ^37^, which integrates the TPMI imputation data and the summary genetic effects from meta-GWASs conducted in various ethnic populations in DIAGRAM ^23^ through a shared continuous shrinkage prior. The populations’ ancestries include: (a) East Asian ancestry: 283,423 individuals (56,268 cases and 227,155 controls); (b) European ancestry: 933,970 Caucasian individuals (80,154 cases and 853,816 controls); (c) South Asian ancestry: 49,492 individuals (16,540 cases and 32,952 controls). Moreover, ancestry-matched linkage disequilibrium references were extracted from the EAS, EUR, and AFR in the 1000 Genomes Project ^17^ – 9,106,250, 10,454,875, and 10,401,621 SNPs for EAS, EUR, and SAS were merged with the TPMI imputation data to calculate the population-specific PRS for each individual using the PLINK (--score command). SNP effect sizes across the three population-specific PRSs were combined using an inverse-variance-weighted meta-analysis of population-specific posterior effect size estimates to calculate a final PRS (--meta command).

In addition, we applied the PGS for T2D (PGS002308) from the PGS Catalog^24^, where the SNP effect sizes were estimated based on 23,827 individuals with African American ancestry, 177,415 individuals with East Asian ancestry, and 898,130 individuals with European ancestry using PRS-CSx ^38^.

The TPMI participants were partitioned into 205,779 independent and 62,304 related participants. Logistic regression models, with T2D disease status as a dichotomous response variable and PRS with and without demographic variables (age and sex) as independent variables, were established based on the independent participants. Finally, the models were applied to the related participants to evaluate the model performance, assessed by AUC.

## Ethics

This study was approved by the Institutional Review Boards of Taipei Veterans General Hospital (2020-08-014A), National Taiwan University Hospital (201912110RINC), Tri-Service General Hospital (2-108-05-038), Chang Gung Memorial Hospital (201901731A3), Taipei Medical University Healthcare System (N202001037), Chung Shan Medical University Hospital (CS19035), Taichung Veterans General Hospital (SF19153A), Changhua Christian Hospital (190713), Kaohsiung Medical University Chung-Ho Memorial Hospital (KMUHIRB-SV(II)-20190059), Hualien Tzu Chi Hospital (IRB108-123-A), Far Eastern Memorial Hospital (110073-F), Ditmanson Medical Foundation Chia-Yi Christian Hospital (IRB2021128), Taipei City Hospital (TCHIRB-10912016), Koo Foundation Sun Yat-Sen Cancer Center (20190823A) Cathay General Hospital (CGH-P110041), Fu Jen Catholic University Hospital (FJUH109001) and Academia Sinica (AS-IRB01-18079), Taiwan. Written informed consent was obtained from the subjects in accordance with institutional requirements and the Declaration of Helsinki principles. All collected information was de-identified before statistical data analysis.

## Acknowledgments

We thank all the participants and investigators from the Taiwan Precision Medicine Initiative. This study was funded by Academia Sinica (40-05-GMM, AS-GC-110-MD02, and 236e-1100202) and National Development Fund, Executive Yuan (NSTC 111-3114-Y-001-001).

## Data availability statement

The genotyping and electronic medical record (EMR) data analyzed in this study are from the Taiwan Precision Medicine Initiative (TPMI) with proper approval from the TPMI Data Access Committee. In compliance with the confidentiality laws governing genetic and health data in Taiwan, the de-identified TPMI data are kept in a secure server at the Academia Sinica and not released to the public. All summary statistics, polygenic risk score (PRS) models, and GWAS results are freely available from the TPMI website (https://tpmi.ibms.sinica.edu.tw). Researchers requesting access to the individual genotyping and EMR data can do so on a collaborative basis. Instructions on requesting access to the data can be found on the TPMI’s official website (https://tpmi.ibms.sinica.edu.tw).

In addition to the TPMI data, we analyzed the following datasets as part of the validation:

- Meta-GWAS summary statistics for T2D across multiple populations from the DIAGRAM Consortium are available at https://diagram-consortium.org/downloads.html.
- The linkage disequilibrium reference from various populations of the 1000 Genomes Project can be downloaded from https://github.com/getian107/PRScsx.
- PRS-CSx weights for T2D across multiple populations from the PGS Catalog are available at https://www.pgscatalog.org/score/PGS002308/.
- The 1000 Genomes Project data can be accessed via PLINK 2.0 at https://www.cog-genomics.org/plink/2.0/resources#phase3_1kg.
- Fastq files of Simons Genome Diversity Project (SGDP) samples are available at https://www.internationalgenome.org/data-portal/data-collection/sgdp.
- Genotype data from the Taiwan Biobank are available through a formal application process (https://www.twbiobank.org.tw/index.php).
- Fastq files of Simons Genome Diversity Project (SGDP) samples are available at https://www.internationalgenome.org/data-portal/data-collection/sgdp.
- Genotype data from the Taiwan Biobank are available through a formal application process (https://www.twbiobank.org.tw/index.php).

In addition, to support researchers in data exploration, TPMI has developed several platforms, including

### TPMI PheWeb

A user-friendly interface that allows researchers to explore associations between genetic variants and phenotypes. It provides access to summary statistics from genome-wide association studies (GWAS) across a broad range of phenotypes and traits. (https://pheweb.ibms.sinica.edu.tw)

### TPMI SNPView

A comprehensive platform that offers detailed information on all genetic variants included in the TPMI SNP arrays, such as Minor Allele Frequency (MAF) among TPMI participants, and variant data from reputable sources like ClinVar, OMIM, and the NCBI dbSNP database. (https://tdap.ibms.sinica.edu.tw/snpview/)

### TPMI DataView

This platform provides statistical insights on TPMI participants, including data on specific health conditions, lab test results, prescribed medications, and treatments received. (https://dataview.ibms.sinica.edu.tw/)

### TPMI Data Analysis Platform (TDAP)

Authorized researchers can access TDAP, a secure central database and analysis platform, once the TPMI Data Access Committee approves their research concepts and their protocols have received approval from Institutional Review Boards (IRBs).

## References

1. Ashley, E.A. Towards precision medicine. Nat Rev Genet 17, 507–522 (2016).

2. Chen, R. & Snyder, M. Promise of personalized omics to precision medicine. Wiley Interdiscip Rev Syst Biol Med 5, 73–82 (2013).

3. Neergard, L. Obama proposes ‘precision medicine’to end one-size-fits-all. Drug Discovery and Development (2015).

4. Landry, L.G., Ali, N., Williams, D.R., Rehm, H.L. & Bonham, V.L. Lack Of Diversity In Genomic Databases Is A Barrier To Translating Precision Medicine Research Into Practice. Health Aff. 37, 780–785 (2018).

5. Popejoy, A.B. & Fullerton, S.M. Genomics is failing on diversity. Nature 538, 161–164 (2016).

6. Wei, C.Y., et al. Genetic profiles of 103,106 individuals in the Taiwan Biobank provide insights into the health and history of Han Chinese. Npj Genom Med 6, 10 (2021).

7. Welter, D., et al. The NHGRI GWAS Catalog, a curated resource of SNP-trait associations. Nucleic. Acids. Res. 42, D1001–D1006 (2014).

8. Richards, S., et al. Standards and guidelines for the interpretation of sequence variants: a joint consensus recommendation of the American College of Medical Genetics and Genomics and the Association for Molecular Pathology. Genet Med 17, 405–424 (2015).

9. Landrum, M.J., et al. ClinVar: public archive of interpretations of clinically relevant variants. Nucleic Acids Res 44, D862–868 (2016).

10. Thorn, C.F., Klein, T.E. & Altman, R.B. PharmGKB summary: very important pharmacogene information for angiotensin-converting enzyme. Pharmacogenet Genomics 20, 143–146 (2010).

11. Hamosh, A., Scott, A.F., Amberger, J.S., Bocchini, C.A. & McKusick, V.A. Online Mendelian Inheritance in Man (OMIM), a knowledgebase of human genes and genetic disorders. Nucleic Acids Res 33, D514–517 (2005).

12. Zhou, W., et al. Efficiently controlling for case-control imbalance and sample relatedness in large-scale genetic association studies. Nat Genet 50, 1335–1341 (2018).

13. Loh, P.R., et al. Efficient Bayesian mixed-model analysis increases association power in large cohorts. Nat Genet 47, 284–290 (2015).

14. Fan, C.T., Lin, J.C. & Lee, C. Taiwan Biobank: A project aiming to aid Taiwan’s transition into a biomedical island. Pharmacogenomics 9, 235–246 (2008).

15. Feng, Y.A., et al. Taiwan Biobank: A rich biomedical research database of the Taiwanese population. Cell Genom 2, 100197 (2022).

16. Mallick, S., et al. The Simons Genome Diversity Project: 300 genomes from 142 diverse populations. Nature 538, 201–206 (2016).

17. Siva, N. 1000 Genomes project. Nat. Biotechnol. 26, 256 (2008).

18. Wang, F.-C. Causes and patterns of ethnic intermarriage among the Hokkien, Hakka, and Mainlanders in postwar Taiwan: a preliminary examination. Bulletin of the Institute of Ethnology, Academia Sinica 76, 43–96 (1993).

19. Liu, D., Ko, A.M. & Stoneking, M. The genomic diversity of Taiwanese Austronesian groups: Implications for the “Into- and Out-of-Taiwan” models. PNAS Nexus 2, pgad122 (2023).

20. Yang, H.C., Chen, C.W., Lin, Y.T. & Chu, S.K. Genetic ancestry plays a central role in population pharmacogenomics. Communications Biology 4(2021).

21. Gauderman, A. QUANTO 1.1: A computer program for power and sample size calculations for genetic-epidemiology studies. http://hydra.usc.edu/gxe (2006).

22. Ministry of Health and Welfare. Statistics of Health Promotion 2021. https://www.hpa.gov.tw/EngPages/Detail.aspx?nodeid=1070&pid=14658 (Health Promotion Administration, 2021).

23. Mahajan, A., et al. Multi-ancestry genetic study of type 2 diabetes highlights the power of diverse populations for discovery and translation. Nat Genet 54, 560–572 (2022).

24. Lambert, S.A., et al. The Polygenic Score Catalog as an open database for reproducibility and systematic evaluation. Nat. Genet. 53, 420–425 (2021).

25. Lin, L.Y., Warren-Gash, C., Smeeth, L. & Chen, P.C. Data resource profile: the National Health Insurance Research Database (NHIRD). Epidemiol Health 40, e2018062 (2018).

26. Hsing, A.W. & Ioannidis, J.P. Nationwide Population Science: Lessons From the Taiwan National Health Insurance Research Database. JAMA Intern Med 175, 1527–1529 (2015).

27. Liu, T.Y., et al. Comparison of multiple imputation algorithms and verification using whole-genome sequencing in the CMUH genetic biobank. Biomedicine-Taiwan 11, 57–65 (2021).

28. Vasimuddin, M., Misra, S., Li, H. & Aluru, S. Efficient architecture-aware acceleration of BWA-MEM for multicore systems. in 2019 IEEE international parallel and distributed processing symposium (IPDPS) 314–324 (IEEE, 2019).

29. Poplin, R., et al. A universal SNP and small-indel variant caller using deep neural networks. Nat Biotechnol 36, 983–987 (2018).

30. Patterson, M., et al. WhatsHap: Weighted Haplotype Assembly for Future-Generation Sequencing Reads. J Comput Biol 22, 498–509 (2015).

31. Delaneau, O., Zagury, J.F., Robinson, M.R., Marchini, J.L. & Dermitzakis, E.T. Accurate, scalable and integrative haplotype estimation. Nature communications 10, 5436 (2019).

32. Hofmeister, R.J., Ribeiro, D.M., Rubinacci, S. & Delaneau, O. Accurate rare variant phasing of whole-genome and whole-exome sequencing data in the UK Biobank. Nat Genet 55, 1243–1249 (2023).

33. Rubinacci, S., Delaneau, O. & Marchini, J. Genotype imputation using the Positional Burrows Wheeler Transform. PLoS Genet 16, e1009049 (2020).

34. Manichaikul, A., et al. Robust relationship inference in genome-wide association studies. Bioinformatics 26, 2867–2873 (2010).

35. R Core Team. R: A language and environment for statistical computing. (R Foundation for Statistical Computing, Vienna, Austria, 2022).

36. Bliss, C. Statistics in biology, (McGraw-Hill, New York, 1967).

37. Ruan, Y., et al. Improving polygenic prediction in ancestrally diverse populations. Nature Genetics 54, 573–580 (2022).

38. Ge, T., et al. Development and validation of a trans-ancestry polygenic risk score for type 2 diabetes in diverse populations. Genome Med 14, 70 (2022).

