## Supplemental file for "The Taiwan Precision Medicine Initiative: A Cohort for Large-Scale Studies"

**Supplemental information of the paper entitled "The Taiwan Precision Medicine Initiative: A Cohort for Large-Scale Studies"**

### Supplemental Information


| | | |
|---|---|---|
| **Figure S1** | TPMI overview and organization structure…………………………………………… | 1 |
| **Figure S2** | Shared SNPs in TPMv1 and TPMv2 arrays………………………………………….. | 2 |
| **Figure S3** | Data types in the TPMI Data Access Platform (TDAP)………………………….. | 3 |
| **Figure S4** | Homozygosity analysis ……………………………………………………………………. | 4 |
| **Figure S5** | Data quality control of samples and SNPs for GWAS……………………………… | 5 |
| **Figure S6** | Quality control check for missing rate, homozygosity rate, and divergent ancestry ……………………………………………………………………………………………… | 6 |
| **Figure S7** | Manhattan plots for a GWAS of Essential Hypertension and two blood pressures ……………………………………………………………………………………………… | 7 |
| **Figure S8** | Manhattan plots for SAIGE analysis of T2D, HbA1c, EHT, and two blood pressures (SBP and DBP) ……………………………………………………………………… | 8 |
| **Figure S9** | Polygenic risk score analysis based on T2D PGS (PGS002308)…………………… | 9 |
| **Table S1** | SNP content of TPMv1 and TPMv2 arrays………………………………………………. | 10 |
| **Table S2** | Electronic medical record data in the TPMI Data Access Platform (TDAP)… | 11 |
| **Table S3** | 83 genetic conditions for Return of Results (ROR)…………………………………… | 12 |

### Supplemental Figures

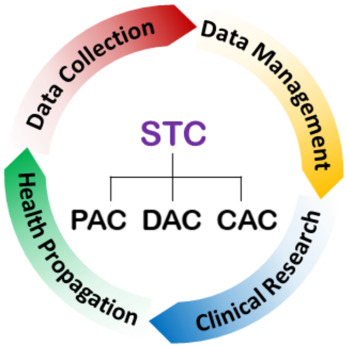


**Figure S1. TPMI overview and organization structure.** The TPMI Consortium is organized by the Steering Committee (STC), the Data Access Committee (DAC), the Clinical Application Committee (CAC), and the Publication Committee (PAC). In the cyclic workflow of TPMI, Data Collection involves enrolling study participants, collecting genetic and EMR data, and developing population-optimized SNP arrays. Data Management includes data quality control and the establishment of the TPMI Data Analysis Platform (TDAP) and Data Lake. Clinical Research encompasses the establishment of working groups, the proposal of research ideas, the development of polygenic risk score (PRS) algorithms, and advanced genetic analyses. Health Propagation involves the return of results (ROR), engaging in public education initiatives, and closely collaborating with non-profit organizations.

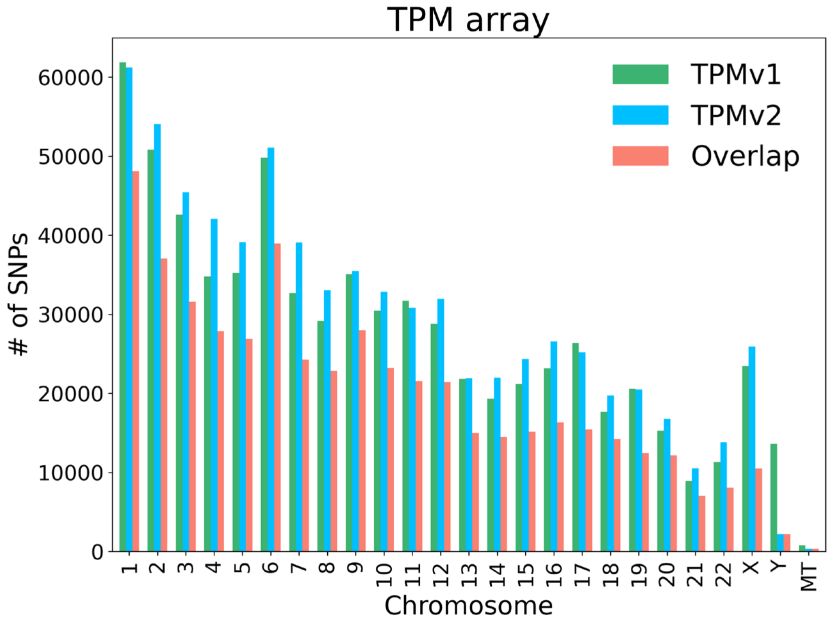


**Figure S2. Shared SNPs in TPMv1 and TPMv2 arrays.** TPMI developed two custom-designed SNP arrays, TPMv1 and TPMv2. TPMv1 interrogates 686,463 SNPs (represented by green bars), and TPMv2 interrogates 743,227 SNPs (represented by blue bars). The two arrays share 494,968 SNPs (represented by red bars), including 481,987 on autosomes, 10,490 on the X chromosome, 2,171 on the Y chromosome, and 320 on the mitochondrial chromosome.

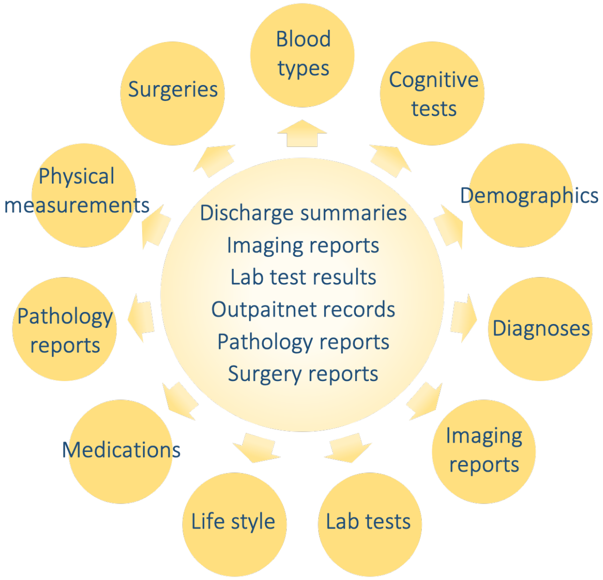


**Figure S3. Data types in the TPMI Data Access Platform (TDAP).** TPMI collected electronic medical records (EMR) data, which includes discharge summaries, imaging reports, lab test results, outpatient records, pathology reports, and surgery reports. Each type of record comprises both free-text sections and predefined structured data. The data can be further divided into the following subcategories: blood types, cognitive tests, demographics, diagnoses, imaging reports, lab tests, lifestyle, medications, pathology reports, physical measurements, and surgeries, totaling more than 140 EMR variables.

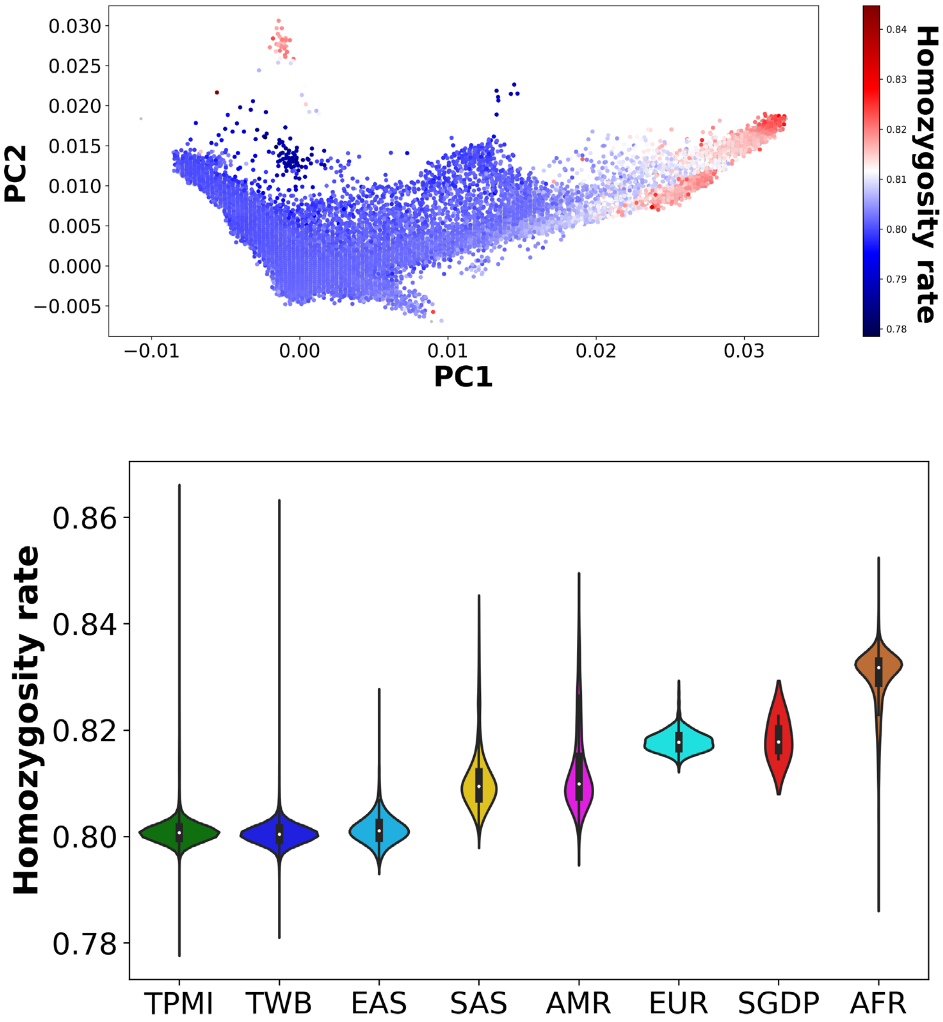


**Figure S4. Homozygosity analysis. (A) Heatmap of homozygosity rate. (B) Violin plots of homozygosity rate.**

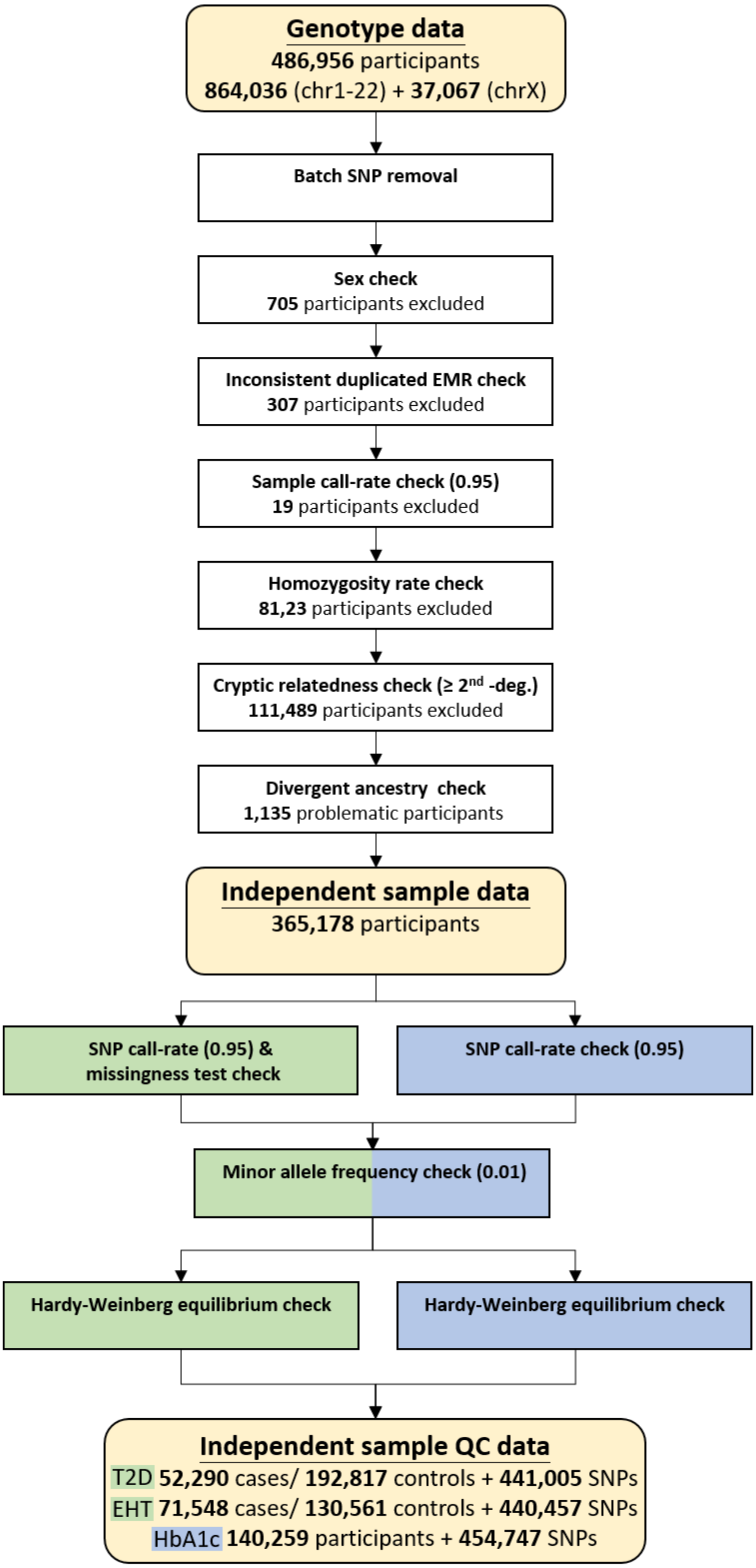


**Figure S5. Data quality control of samples and SNPs for GWAS.**

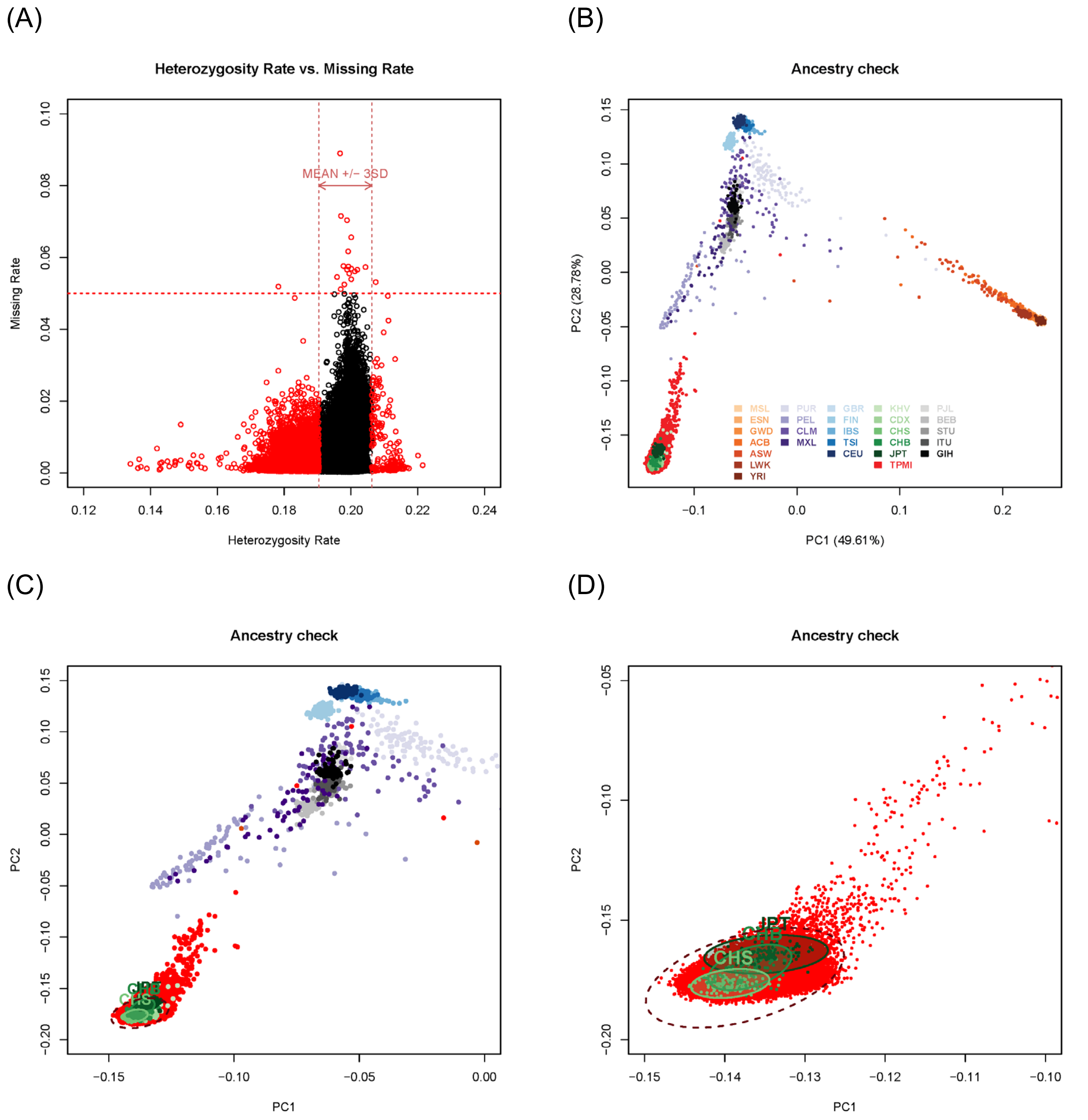


**Figure S6. Quality control check for missing rate, homozygosity rate, and divergent ancestry. (A) Quality control check for missing rate and homozygosity rate. (B) Ancestry check using PCA for global populations. (C) Ancestry check using PCA for non-African populations. (D) Ancestry check using PCA for Asian populations.**

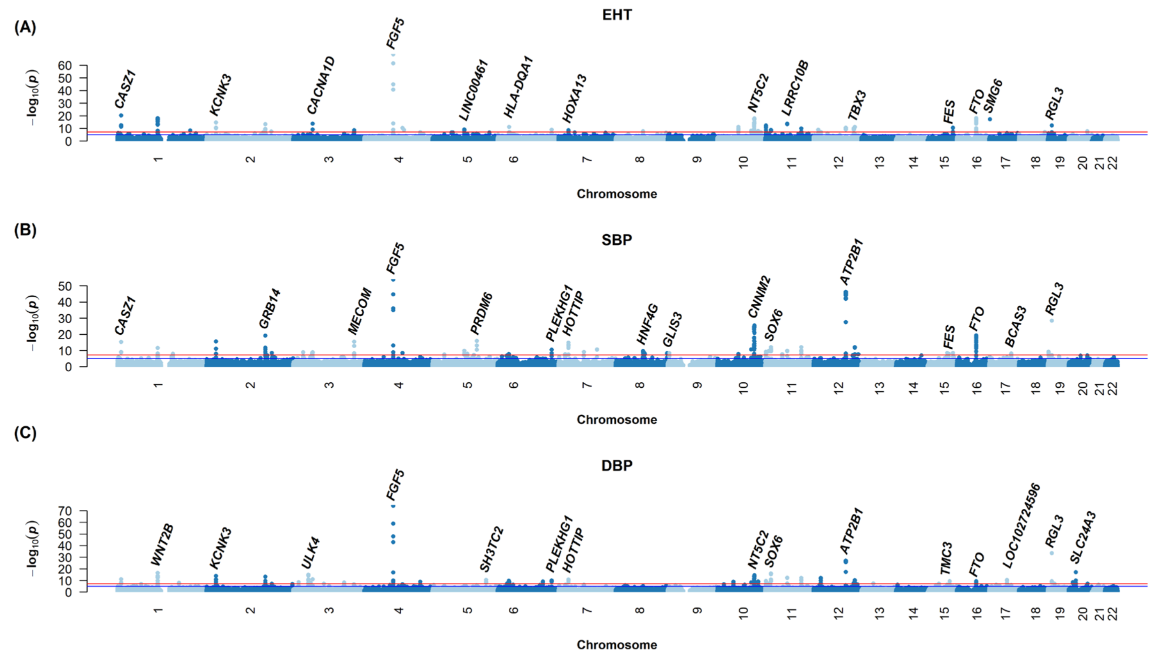


**Figure S7. Manhattan plots for a GWAS of Essential Hypertension and two blood pressures. (A) Essential Hypertension (EHT, 71,548 cases and 130,561 controls); (B) Systolic blood pressure (SBP, n = 241,667); (C) Diastolic blood pressure (DBP, n = 241,646).**

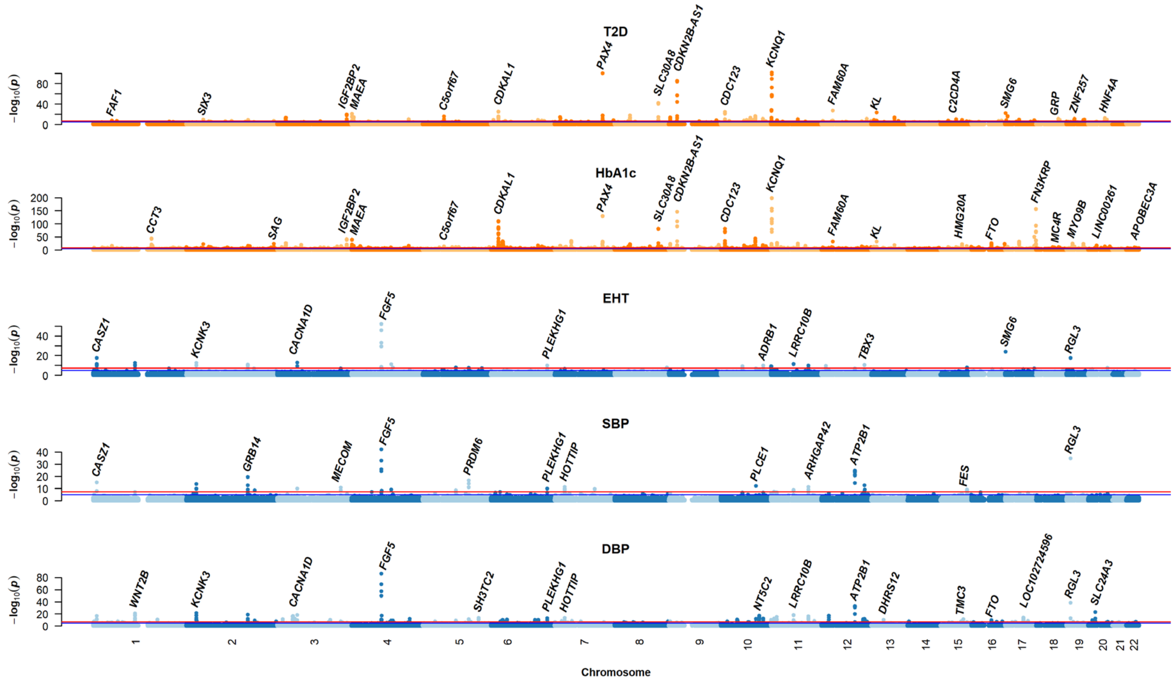


**Figure S8. Manhattan plots for SAIGE analysis of T2D, HbA1c, EHT, and two blood pressures (SBP and DBP). (A) GWAS of Type 2 Diabetes (T2D) (n = 52,290 cases and 192,817 controls); (B) QTL mapping of Hemoglobin A1c (HbA1c) (n = 140,259); (C) GWAS of Essential Hypertension (EHT) (n = 71,548 cases and 130,561 controls); (D) QTL mapping of systolic blood pressure (SBP) (n = 241,667); (E). QTL mapping of diastolic blood pressure (DBP) (n = 241,646).**

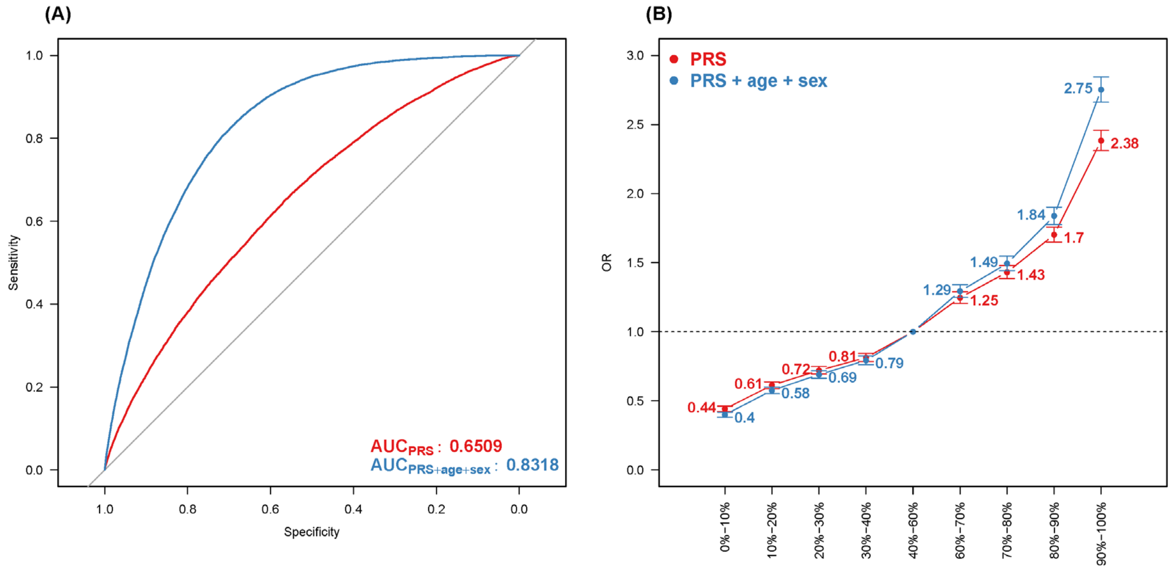


**Figure S9. Polygenic risk score analysis based on T2D PGS (PGS002308). (A) Area under the receiver operating characteristic curve (AUC)**. AUC of PRS (red curve) and AUC of PRS, age, and sex (blue curve) are displayed. **(B) Dose-response effect of PRS levels on the odds ratio of T2D**. Dose-response effect of PRS levels on the odds ratio of T2D (red line) and dose-response effect of a combination of PRS, age and sex (blue curve).

### Supplemental Tables

**Table S1. SNP content of TPMv1 and TPMv2 arrays.** TPMv1 contains 686,463 SNPs, and TPMv2 contains 743,227 SNPs. Note that some SNPs from different categories may overlap.

| Category | TPMv1 | TPMv2 |
|---|---|---|
| GWAS Grid | 403,082 | 580,364 |
| GWAS Catalog | 67,002 | 41,645 |
| ACMG | 40,295 | 2,803 |
| ClinVar | 90,343 | 15,467 |
| PharmGKB | 3,949 | 2,911 |
| OMIM | 82,425 | 165,631 |
| Loss of function (LoF) | 49,794 | 48,477 |
| Copy number variation (CNV) | 42,660 | 22,884 |

GWAS Catalog: https://www.ebi.ac.uk/gwas/
ACMG: https://www.acmg.net/
ClinVar: https://www.ncbi.nlm.nih.gov/clinvar/
PharmGKB: https://www.pharmgkb.org/
OMIM: https://www.omim.org/

**Table S2. Electronic medical record data in the TPMI Data Access Platform (TDAP).**

| Record type | Outpatient records | Discharge summary | Lab test results | Pathology reports | Surgery reports | Imaging reports |
|---|---|---|---|---|---|---|
| Free text data | Division, Condition summary, Family history | Division, Chief complaint, Laboratory data, Imaging study, Hospital course, Complications | Reference range, Remarks | Reference range, Remarks | Adverse drug, Blood Transfusion reaction, history of transplantation | Division, Findings, Chief complaint, Results |
| Structured data | Age, Sex, Date, Blood type, Major illness, ICD diagnosis, Procedures, Prescription | Age, Sex, Date, ICD diagnosis, Cancer staging | Age, Sex, Date, NHI code, test results | Age, Sex, Date, Major illness, ICD diagnosis, Specimen site, Result | Age, Sex, Date, Blood type, Pre- / Post-operation ICD diagnosis, Surgical approach, Operative Finding | Age, Sex, Date, Image type, Body areas, ICD diagnosis |

ICD diagnosis is based on ICD-9 or ICD-10.

**Table S3. 83 genetic conditions for Return of Result (ROR).** (A) Genetic conditions for health management; (B) Genetic conditions for disease diagnosis; (C) Pharmacogenetic associations for medicines.

| | **Disease / Medicine** | **Gene** |
|---|---|---|
| Genetic conditions for health management | Alcohol flushing reaction | *ALDH2* |
| | Hereditary cancer-predisposing_syndrome/Breast-ovarian cancer, familial 4 | *RAD51D* |
| | Familial hypercholesterolemia | *APOB, LDLR* |
| | DFNA 2 Nonsyndromic Hearing Loss | *KCNQ4* |
| | Hereditary pancreatitis | *PRSS1, SPINK1* |
| | Cerebral autosomal dominant arteriopathy with subcortical infarcts and leukoencephalopathy (CADASIL) | *NOTCH3* |
| | Hyperuricemia | *ABCG2* |
| Genetic conditions for disease diagnosis | Glucose-6-phosphate dehydrogenase deficiency | *G6PD* |
| | Meckel syndrome | *MKS1* |
| | beta Thalassemia | *HBB* |
| | Total iodide organification defect/Neonatal transient hypothyroidism | *TPO* |
| | congenital hypothyroidism | *TSHR* |
| | PMM2-congenital disorder of glycosylation | *PMM2* |
| | primary carnitine deficiency | *SLC22A5* |
| | Wilson disease | *ATP7B* |
| | Joubert Syndrome | *CEP290* |
| | primary autosomal recessive microcephaly 1 | *MCPH1* |
| | glutaric acidemia type I | *GCDH* |
| | Krabbe Disease (Later-onset/adult-onset form) | *GALC* |
| | citrin deficiency (adult-onset type II citrullinemia) | *SLC25A13* |
| | sialidosis | *NEU1* |
| | Waardenburg syndrome | *EDNRB* |
| | glycogen storage disease type 1A | *G6PC* |
| | limb-girdle muscular dystrophy-7 (LGMDR7) | *TCAP* |
| | limb-girdle muscular dystrophy-18 (LGMDR18) | *TRAPPC11* |
| | cystic fibrosis | *CFTR* |
| | phenylketonuria | *PAH* |
| | 6-pyruvoyltetrahydropterin synthase deficiency | *PTS* |
| | MYH-associated polyposis | *MUTYH* |
| | hereditary factor XI deficiency disease | *F11* |
| | hereditary spastic paraplegia type 5 | *CYP7B1* |
| | hereditary spastic paraplegia type 15 | *ZFYVE26* |
| | ABCA4-Related Disorders (cone-rod dystrophy/Stargardt macular degeneration) | *ABCA4* |
| | autosomal recessive bestrophinopathy | *BEST1* |
| | Cone-rod dystrophy and hearing loss 1 (CRDHL1) | *CEP78* |
| | Nonsyndromic hearing loss and deafness | *GJB2* |
| | Pendred syndrome | *SLC26A4* |
| | Nagashima-type palmoplantar keratosis (NPPK) | *SERPINB7* |

| | | |
|---|---|---|
| | mucolipidosis type III | *GNPTAB* |
| | glycogen storage disease type II | *GAA* |
| Pharmacogenetic associations for drug safety | Lidocaine and Prilocaine | *G6PD* |
| | Ropivacaine | *G6PD* |
| | Celecoxib | *CYP2C9* |
| | Flurbiprofen | *CYP2C9* |
| | Ibuprofen | *CYP2C9* |
| | lornoxicam | *CYP2C9* |
| | meloxicam | *CYP2C9* |
| | Piroxicam | *CYP2C9* |
| | Sulfasalazine | *G6PD, NAT2* |
| | Clopidogrel | *CYP2C19* |
| | Simvastatin | *SLCO1B1* |
| | Glimepiride | *G6PD* |
| | Glipizide | *G6PD* |
| | Rasburicase | *G6PD* |
| | Amikacin | *MT-RNR1* |
| | Ceftriaxone | *G6PD* |
| | Gentamicin | *MT-RNR1* |
| | Hydroxychloroquine | *G6PD* |
| | Isoniazid | *NAT2* |
| | Efavirenz | *CYP2B6* |
| | Nalidixic Acid | *G6PD* |
| | Neomycin | *MT-RNR1* |
| | Paromomycin | *MT-RNR1* |
| | Peginterferon Alfa-2a | *IFNL3* |
| | Peginterferon Alfa-2b | *IFNL3* |
| | Ribavirin | *IFNL3* |
| | Sulfamethoxazole and Trimethoprim | *G6PD, NAT2* |
| | Streptomycin | *MT-RNR1* |
| | Tobramycin | *MT-RNR1* |
| | Carisoprodol | *CYP2C19* |
| | Flutamide | *G6PD* |
| | Irinotecan | *UGT1A1* |
| | Nilotinib | *UGT1A1* |
| | Mercaptopurine | *TPMT* |
| | Pazopanib | *UGT1A1* |
| | Citalopram | *CYP2C19* |
| | Escitalopram | *CYP2C19* |
| | Sertraline | *CYP2C19* |
| | Clobazam | *CYP2C19* |
| | Azathioprine | *TPMT* |
| | Tacrolimus | *CYP3A5* |
| | Methylene Blue | *G6PD* |
| | Sodium Nitrite | *G6PD* |
